# A widely applicable and cost-effective method for general and specific RNA-protein complex isolation

**DOI:** 10.1101/2022.03.28.486031

**Authors:** Sam Balzarini, Roosje Van Ende, Arnout Voet, Koen Geuten

## Abstract

Despite important methodological advances made in the past few years, a widely applicable, cost-effective and easily scalable procedure that can be routinely used to isolate ribonucleoprotein complexes (RNPs) remains elusive. We describe a versatile method that connects aspects of existing methods in a workflow optimized to reach the above goals and called it “Silica-based Acidic Phase Separation (SAPS)-capture”. To validate the method, the 18S rRNP of *S. cerevisiae* was captured. To illustrate its applicability, we isolated a repertoire of RNPs in *A. thaliana*. This procedure can provide the community with a powerful tool to advance the study of ribonomes and RNPs in any organism or tissue type.

## BACKGROUND

The interplay between proteins and RNA (the ribonome) plays an important functional role in cell biology. Some important processes regulated by conventional RNA-binding proteins (RBPs), such as the regulation of translation and post-translational modifications (1), have been known for decades. However, recent proteome-wide studies revealed hundreds of novel RBPs without classical RNA-binding domains and raised the concept of not only proteins regulating RNA but as well the potential of RNA to regulate protein function (2). New functions can be attributed to the dynamics of RNA-protein complex (RNPs) formation: the formation of RNP bodies (e.g stress granules) driven by liquid-liquid phase separation (3), the potential of long noncoding RNA to scaffold protein complexes (4), the role of aberrant RBPs in certain diseases (5) amongst others. With the emerging understanding of the importance of these complexes, the impetus to develop novel techniques to study RNPs increased. RIC was the first RNA-centric method to allow the isolation of the mRNA interactome targeting the RNPs poly-A tail (6),(7). Multiple other techniques such as CARIC (8), RICK (9), TRAPP (10), VIR-CLASP (11) to isolate a compilation of RNPs have been developed since. Methods to isolate specific RNP complexes also emerged such as ChiRP-MS (12), CHART-MS (13) and RAP-MS (14). Recently, new methods, based on the well-known organic phase separation (15)(16)(17), were developed to isolate a compendium of RNPs without a bias towards certain RNA sequence elements or post-translational modifications (for an extensive review of these methods see Van Ende et. al. 2020 (18)). Despite the set of techniques currently available, to our knowledge only a few interactomes of single RNA species have been identified (12),(14),(19),(20),(21),(22),(23),(24),(25)..

We believe that technical and cost limitations of previous procedures prevent methods from being routinely applied: (**1**) The complex nature of cell lysate makes downstream applications, such as probe capture, suboptimal. The presence of RNases/proteinases/secondary metabolites requires harsh denaturing buffers and this affects the probe binding specificity so that LNA probes or probe TILING approaches become required (26)(12)(14). These approaches also make the procedures more costly. (**2**) The limited scalability of protocols makes it costly to reach sensitivity thresholds. If scalable, such as the organic phase separation methods or TRAPP, the product is not necessarily free from contaminating RNA, DNA or proteins, which is again suboptimal for downstream applications. (**3**) Limited sensitivity has also been tackled by protein labelling strategies, such as SILAC labelling or PAR cross-linking, but these methods are restricted to cell cultures/systems. (**4**) Multilayer tissues do not allow UV light to penetrate deeply to effectively cross-link RNA and proteins into complexes. This can be further limited by, for example, the presence of a cell wall or UV absorbing molecules. The common method for cross-linking multilayer tissues is currently formaldehyde because it can penetrate these tissues better. This cross-linking type was recently used for a recent RNA-targeting protocol succeeded in isolating a specific mRNA from plant tissue (25). Yet UV cross-linking provides a very specific cross-link between RNA and proteins and is therefore preferable (18). (**5**) Transgenic tissues allow for specific isolation procedures based on sequence tags. Yet such tagging approaches (eg. Ribotrap(27), TRAP(28), MS2-biotrap(29) etc.) modify the wild-type sequence and therefore could modify the result. In addition, they take more time.

Therefore, a definite need exists to integrate aspects of available procedures and establish a broadly applicable protocol to cost-effectively isolate either the interactome (RBPome) of a tissue or a defined interactome for a specific RNA molecule of interest. Here, we believe we established such a strategy by optimizing a combination of key steps from previously described methods. The protocol first pre-isolates all UV cross-linked RNPs, the so-called RBPome, by combining a silica-based purification of RNA and RNPs, subsequently followed by an AGPC (acid guanidinium thiocyanate-phenol-chloroform) extraction, further separating the RNP complexes from free RNA. We refer to this as silica-based acidic phase separation or SAPS. An important advantage of this combination is that it better purifies RNPs from contaminating RNA and proteins and that it does not specifically select RNA at this stage, such as in RIC which selects based on poly-A tailed sequences (6),(7). The SAPS isolate can be used to study the dynamics of the RNA-binding proteome in different environments and biological cues. In our study, the SAPS RBPome was used as an improved starting point to capture a specific RNP of interest (hence SAPS-capture). To validate the SAPS-capture protocol, we isolated the well-characterized 18S rRNP of *Saccharomyces cerevisiae*. This complex was chosen because **(I)** the cryoEM structures of the ribosomal complex and its known 33 protein interactors are described elaborately in the literature (30), and **(II)** previous RNA-targeting protocols, such as capture with the use of LNA/DNA mixmer probes (26) and RAP-MS (14) have also used this complex as a validation of their protocol, enabling a comparison of approaches. While the 18S rRNP is an abundant complex and therefore does not allow to demonstrate sensitivity, it indicates the potential of the method to be immediately applied to other rather abundant targets for example the capture of viral RNPs. The application to lower abundant targets is revised in the discussion. Next, we applied the SAPS protocol to whole tissue samples of *Arabidopsis thaliana*, previously described (31) as difficult to process. Once a successful SAPS is established for such a sample, the RBPome loses its tissue type-dependent character and allows for a universal and streamlined continuation of an RNP capture procedure.

## RESULTS

### Experimental strategy to generalize the procedure

Figure 1 represents a schematic overview of both the pre-RNP isolation (SAPS) and specific RNP isolation procedure (SAPS-Capture).

**Figure 1.**
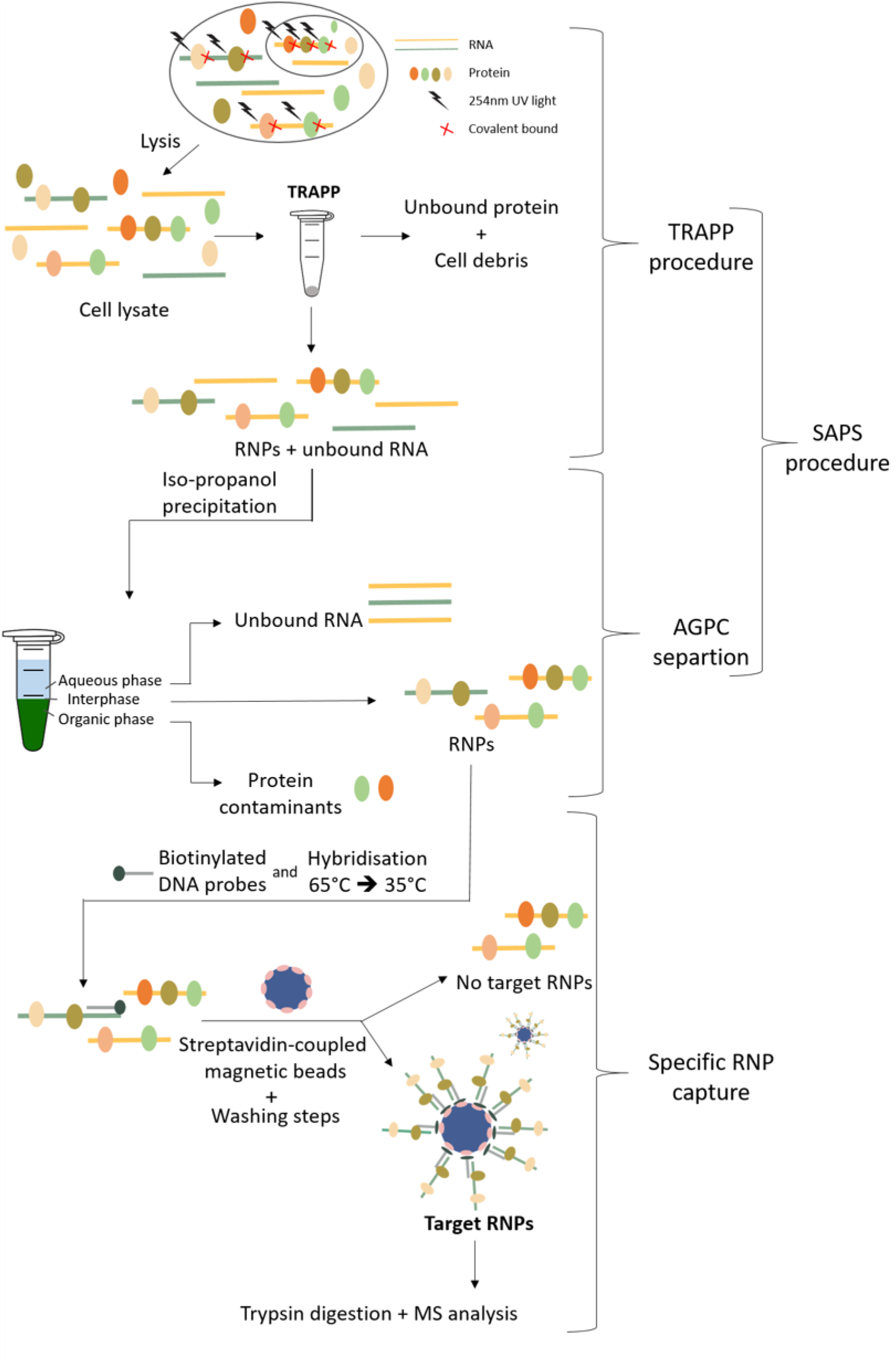
Schematic of the SAPS protocol combined with specific RNP isolation workflow.

### Silica-based acidic phase separation (SAPS)

The SAPS procedure can be divided into a solid phase (adapted TRAPP) and AGPC liquid phase separation.

#### Silica pre-purification of unbound RNA and RNP complexes

In a first step, both unbound RNA and RNPs were purified according to the protocol (total RNA-associated protein purification (TRAPP)) described by Shchepachev et al. (2019) (10) with minor modifications. The TRAPP protocol isolates cross-linked RNP complexes and unbound RNA molecules purifying the mixture from most unbound (also naturally biotinylated) proteins, gDNA, lipids and other macromolecules.

#### DNase treatment

Although both the washing steps and the acidic pH of the TRAPP protocol are designed to reduce the recovery of DNA (32), it is shown that DNA is still partially present in the eluate. In the subsequent AGPC separation, the neutral DNA molecules will dissolve in the organic phase. However, after the TRAPP protocol, designed for processing large quantities of material, the sample is concentrated with isopropanol precipitation and further processed in only small volumes of TRIsure^TM^. Therefore, the recovery of DNA after TRAPP is concentrated as well and can cause saturation of the organic phase with settling on the interphase as a result.

To investigate the presence of DNA, we treated the RNPs after TRAPP with RNase and Benzonase, the latter degrades both RNA and DNA. While RNase-based degradation was rather partial resulting in a smear and RNP complexes getting stuck in the gel pocket, the Benzonase cleavage was complete, showing clear protein bands and an empty gel pocket (Additional file 1). This could be due to the more general cleaving nature of Benzonase compared to RNase, suggesting the presence of DNA after SAPS. Alternatively, this result could also be explained by the pre-heating of the sample when treating with Benzonase, potentially reversing liquid-liquid phase separation of the complexes making them better accessible to the enzyme. More straightforward evidence for the presence of DNA remnants after TRAPP is the high qPCR signal for a non-reverse-transcribed sample. However, due to the low efficient UV cross-linking of proteins to DNA at 254nm, the recovery of DNA-bound proteins is negligible (10). For this reason, a DNase treatment is not required when only the dynamics of the RBPome is studied (could be included for reducing sample complexity however). On the other hand, for the samples prepared for specific RNA-capture, the DNA would occupy the probes and thereby decrease the RNP/bead ratio and a DNase treatment is required.

After DNase treatment, qPCR analysis of the non-reverse-transcribed sample approached the values of the non-template control. In *S. cerevisiae* 25S, as well as 18S, are located on chromosome XII (nucleosome). With 18S DNA present theoretically 25S DNA is captured as well. With the DNase treatment included, we noticed indeed a reduced amount of 25S rRNA/DNA contamination in the 18S rRNA interactome (Figure 2).

**Figure 2.**
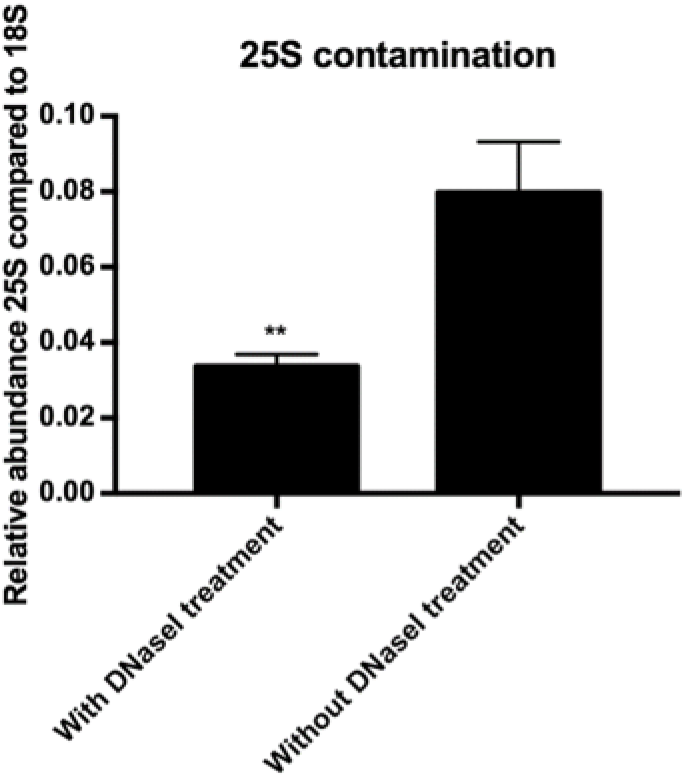
25S rRNA abundance after capture with 18S probes comparing with and without DNase treatment. ** represents a two-tailed p-value<0.01. Error bars represent SEM, n=2.

#### Isopropanol precipitation and AGPC isolation of RNP complexes

After TRAPP, although stringent washing conditions and chaotropic reagents were used, a low level of protein recovery is described in the non-cross-linked samples (10). We perform an AGPC isolation to deplete these protein contaminants. Importantly not only proteins remaining after TRAPP are removed, but also the majority of unbound RNA molecules will be depleted resulting in a far more efficient RNA-capture protocol, as described by Van Ende et al. (2020) (18). Unbound proteins will settle in the organic phase, unbound RNA will settle in the aqueous phase and RNPs, complexes harbouring both hydrophobic and hydrophilic properties, as was recently shown (16),(17),(15), will settle on the interphase (Figure 3). After isolation of the interphase, the RNPs are dissolved in a general buffer of choice. This flexibility of buffers enables downstream applications such as specific RNP isolation to be organism and sample independent.

**Figure 3.**
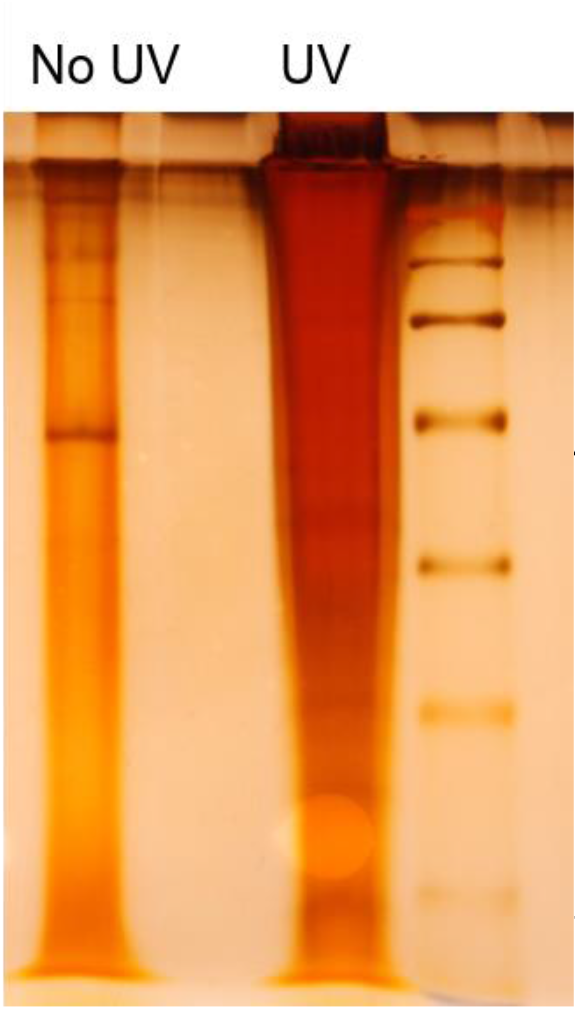
Comparison of non-cross-linked and UV cross-linked sample after SAPS.

### The capture of a specific RNA of interest

After the SAPS purification of RNA-protein complexes, the RNPs are dissolved in a buffer of choice, compared to the frequently used cell lysates to capture on (14),(12),(13). The sample stability due to the absence and denaturation of RNases and proteinases during the purification enables this buffer flexibility. Therefore, subsequent downstream procedures, such as specific RNP capture, can be optimized without a sample-type-origin dependence. The isolated RNPs after the SAPS purification provide a uniform/organism independent input sample for specific RNP capture. This sample is not only depleted of unbound RNA competing with the complexes for probe hybridization but will be as well free from naturally biotinylated molecules, hybridizing with the streptavidin-coated magnetic beads, avoiding an expensive preclearance step.

To validate the SAPS-capture protocol the well-characterized 18S rRNP of *S. cerevisiae* was isolated.

### Quality control (RT-qPCR and BioAnalyzer)

For the detailed outline of the procedure see methods.

The purity of the SAPS-capture samples was checked with RT-qPCR. Often the validation of this type of protocol is presented as a fold-enrichment. This is a comparison between the ratio of the target RNA and a reference gene before and after the experiment. However, it is possible that in absolute numbers the target RNA does not exceed the reference gene, but it is enriched manifold (18). We compared the abundance of both 25S and taf10 with the abundance of 18S only after the experiment. For every 18S molecule only 0.03 25S molecules are present, for taf10 this is only 1E-6molecules (Figure 4A). The amount of 18S rRNA captured with the probes specifically targeting this molecule and the scrambled probes was compared. There appeared to be 6% 18S rRNA present in the control sample compared to the sample with the specific probes (Figure 4B). This data indicates the highly specific nature of the protocol, which is also visually observed. The beads loaded with the ‘empty’ scrambled probes form a dense structure against the magnet, the beads loaded with the 18S probes on the other hand form a more diffuse structure (Additional file 2). Interestingly, it is noticeable that the yield determined as the absolute copy number of 18S molecules after capture increased compared to the input samples. We believe a more efficient reverse transcriptase reaction occurs in the samples after the capture because no competition from other RNA molecules is present. Alternatively, the heating of the sample to elute the complexes might reverse liquid-liquid phase separation resulting in a more efficient reverse transcriptase reaction compared to the non-heated sample before capture.

**Figure 4.**
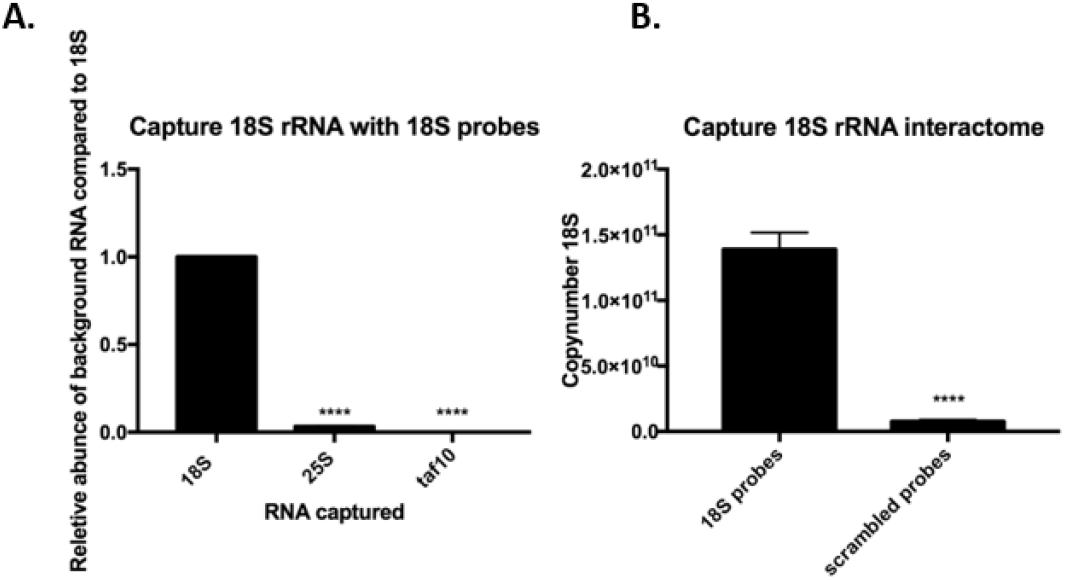
Quantification of the contamination after capturing 18S. **** represents a two-tailed p-value<0.0001 A. Relative abundance of 25S rRNA and taf10 compared to 18S rRNA after capture with 18S probes. Values are calculated as the amount of background after capture divided by the amount of 18S after capture. Error bars represent SEM, n=2. Significance is determined with an unpaired t-test. B. Yield of 18S rRNA comparing captured with 18S probes with capture with scrambled probes. Values are calculated by extrapolation on a standard curve of the plasmid PGEM-T_fulllength18S. Error bars represent

The purity of the samples is again confirmed by BioAnalyzer. Here, the integrity is partially affected potentially due to UV cross-linking or the elution at 95 degrees (Figure 5). However, this is circumvented in the final protocol by an on-bead trypsin digestion. It is shown that after the RNA-capture experiment, the target 18S rRNA is strongly enriched and the other abundant ribosomal RNAs (5S/5.8S and 25S) are depleted (Figure 5B). The control sample is as well completely depleted (Figure 5C).

**Figure 5.**
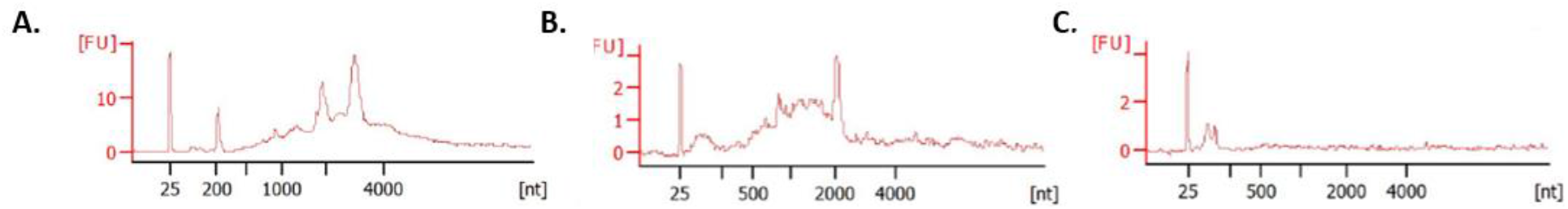
RNA pico BioAnalyzer (Agilent) A. SAPS purification B. Capture with 18S probes C. capture with scrambled probes.

### LC-MS/MS analysis 18S rRNA interactome

A set of 54 proteins (Table 1) was significantly enriched either quantitatively (adjusted p-value <0.01; |log2FC|>2) or semi-quantitatively (no value for the scrambled probes and a value for at least four out of five replicates for the 18S probes) for the 18S rRNA interactome when comparing with a capture with scrambled probes (Figure 6).

**Table 1.**
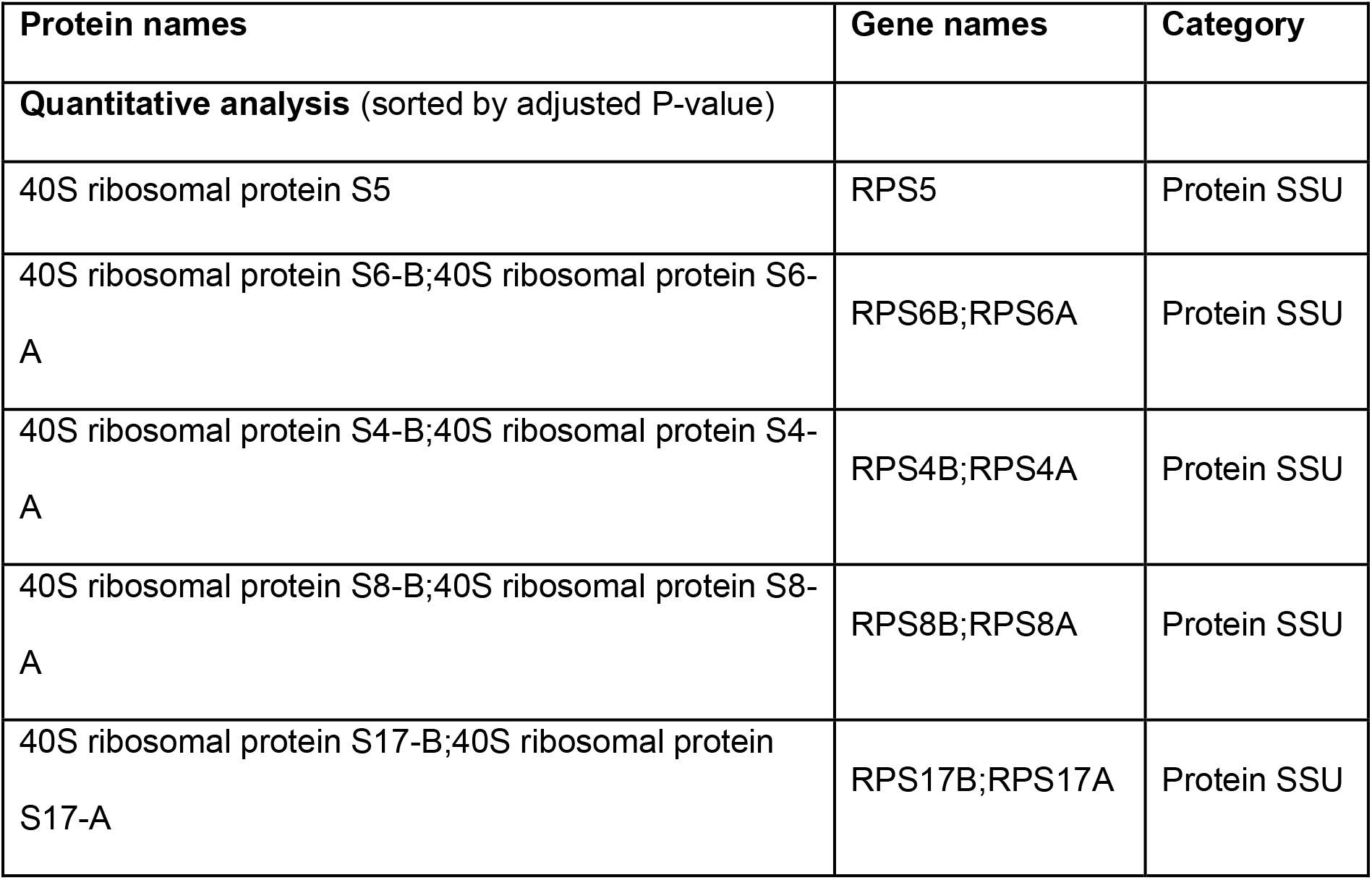
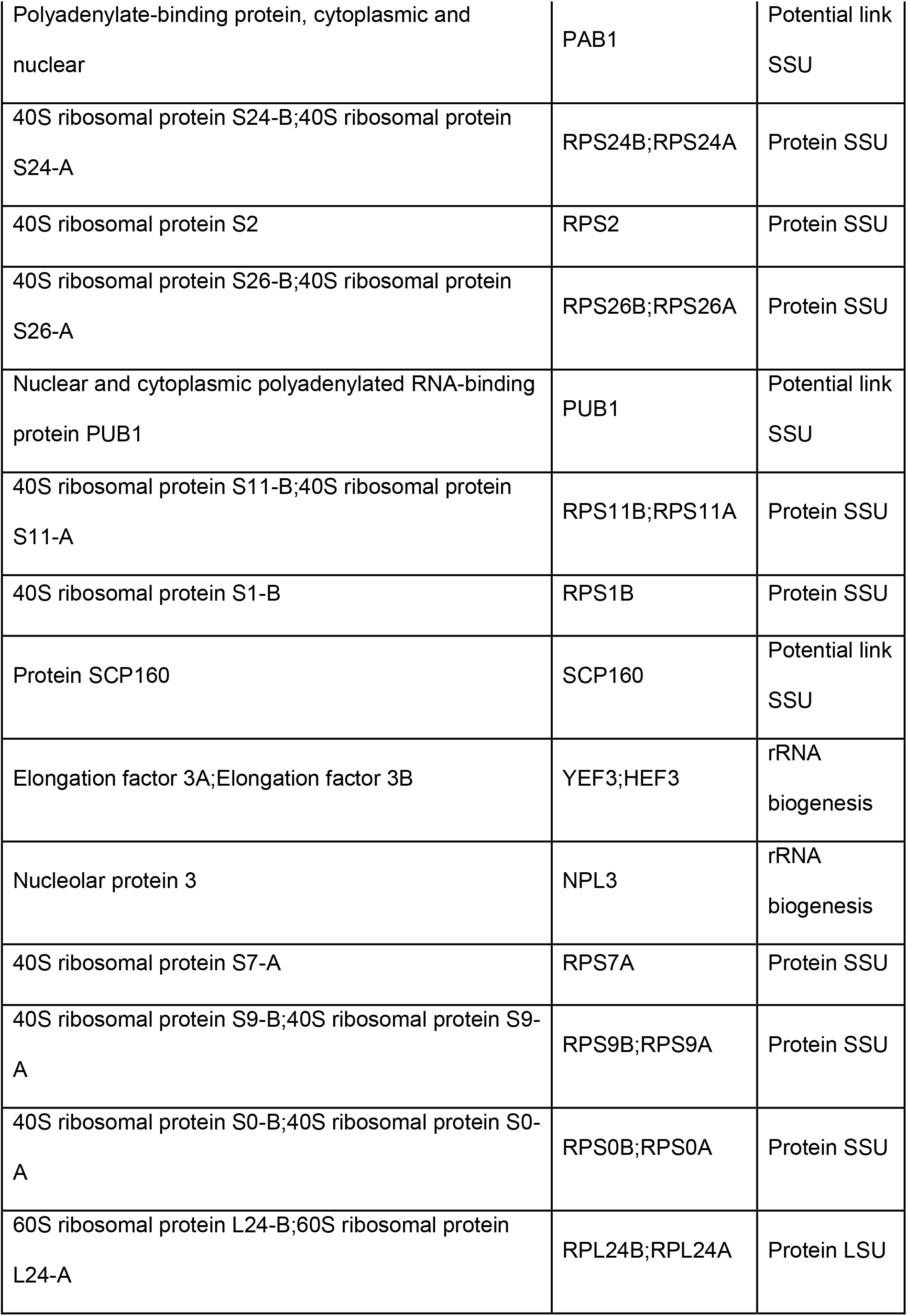
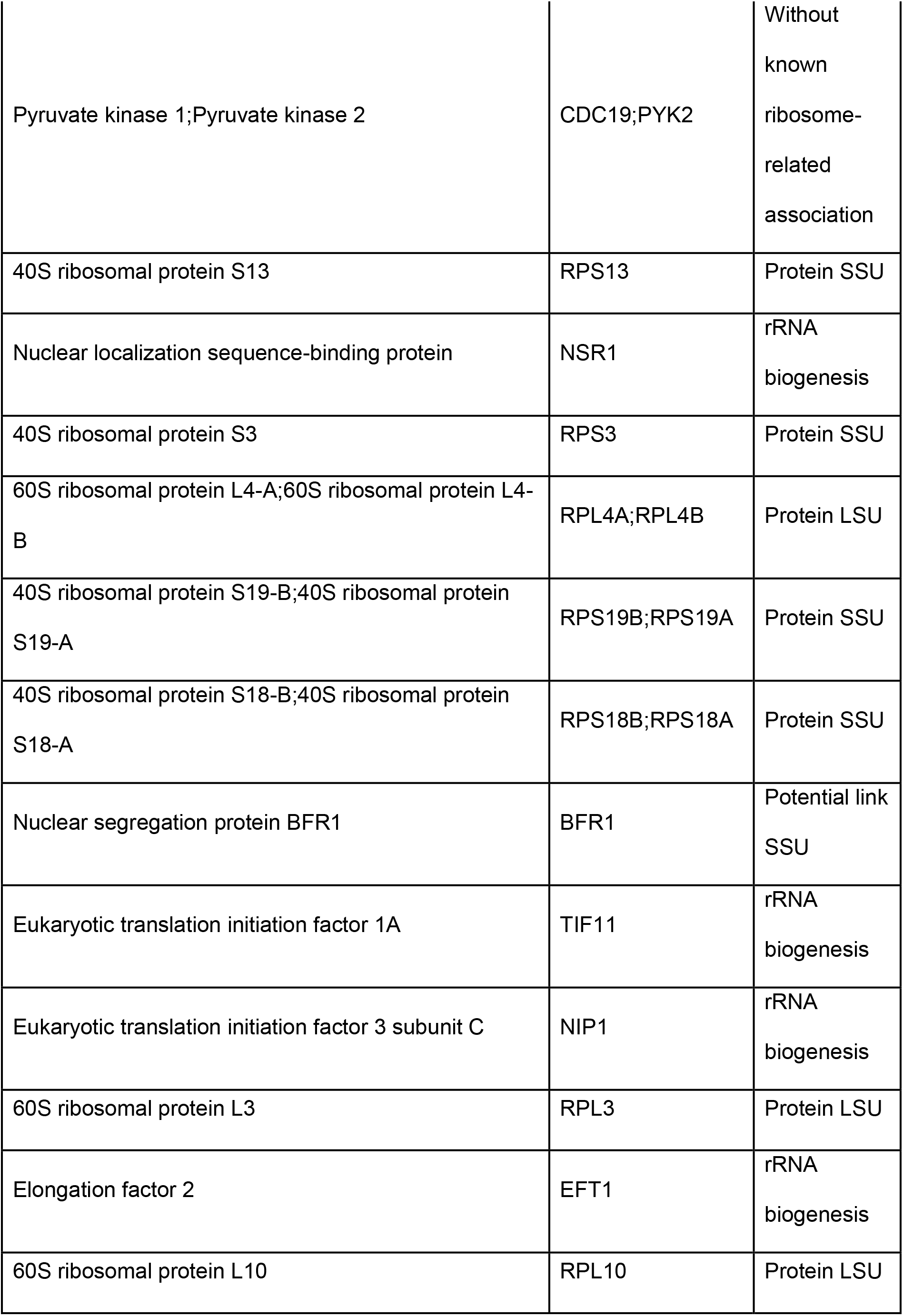
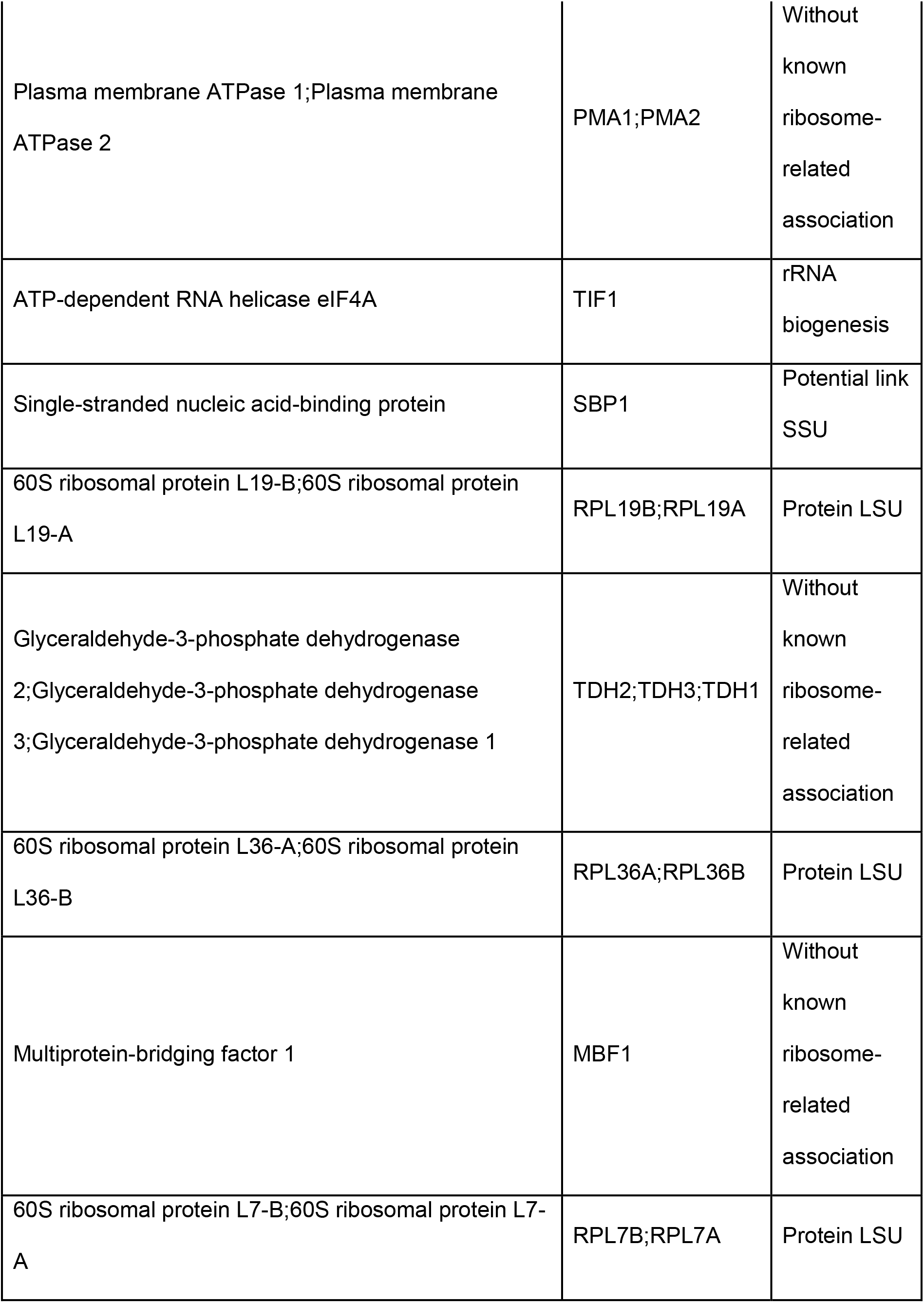
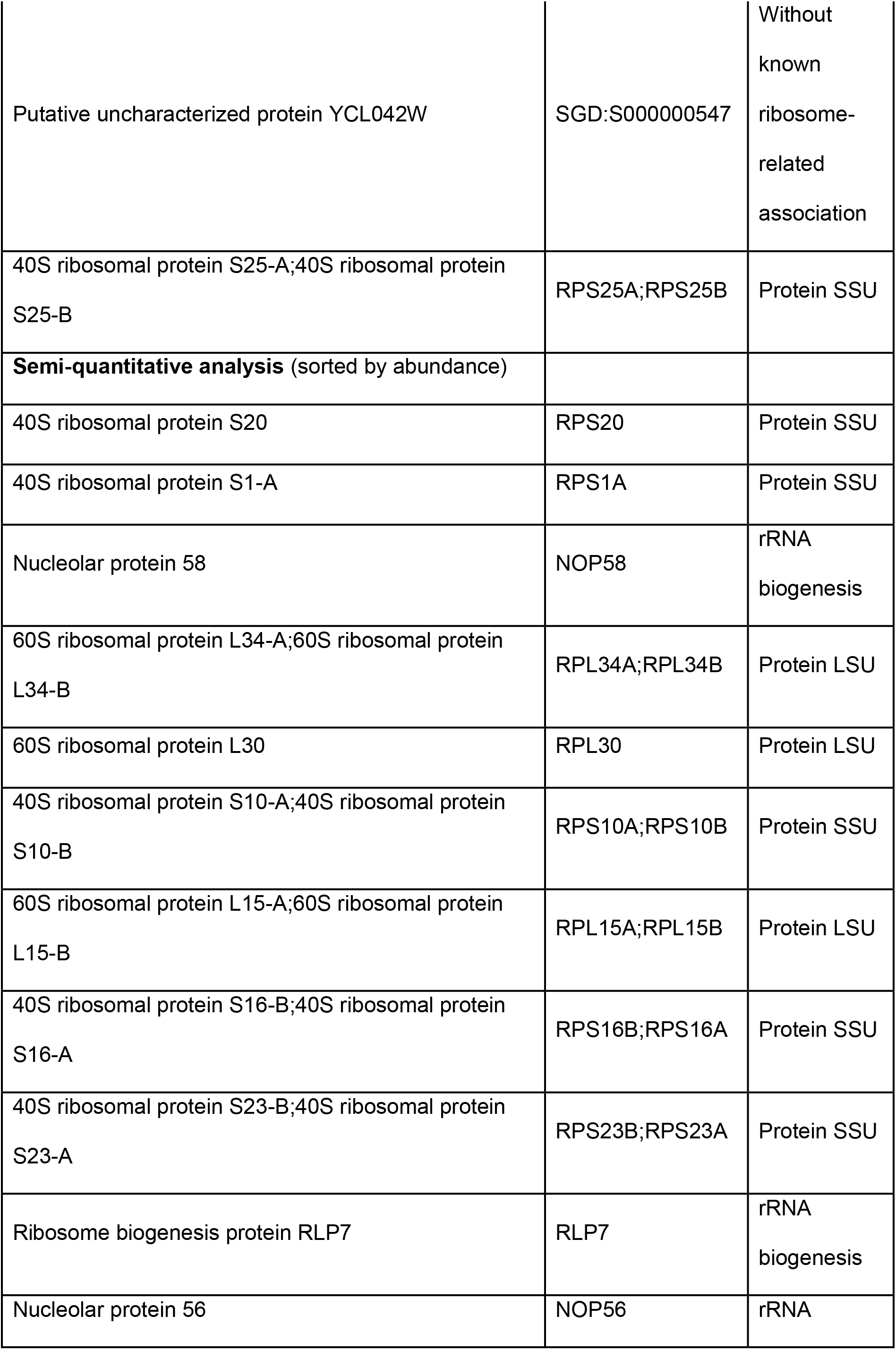
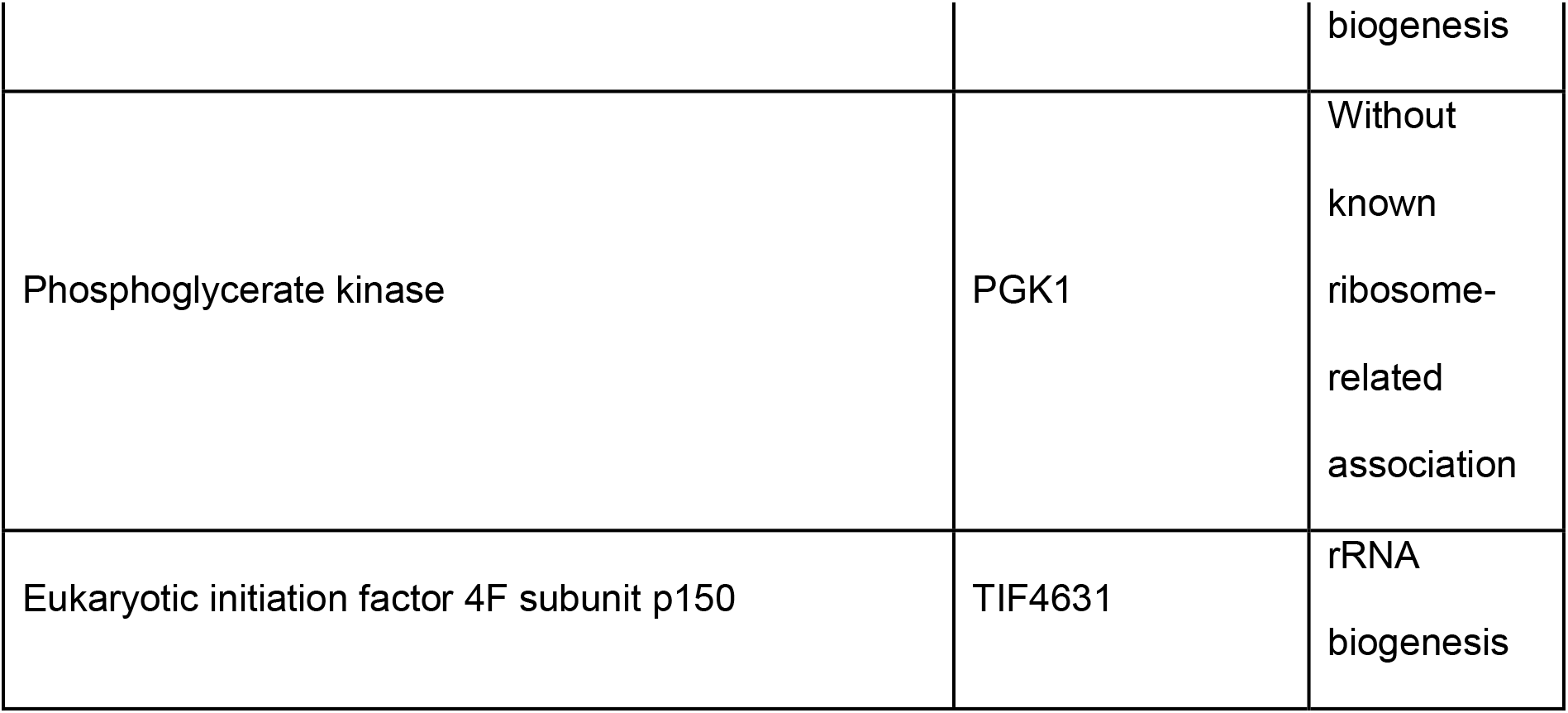
. significantly enriched protein of 18S rRNA interactome (small subunit=SSU, large subunit=LSU)

**Figure 6.**
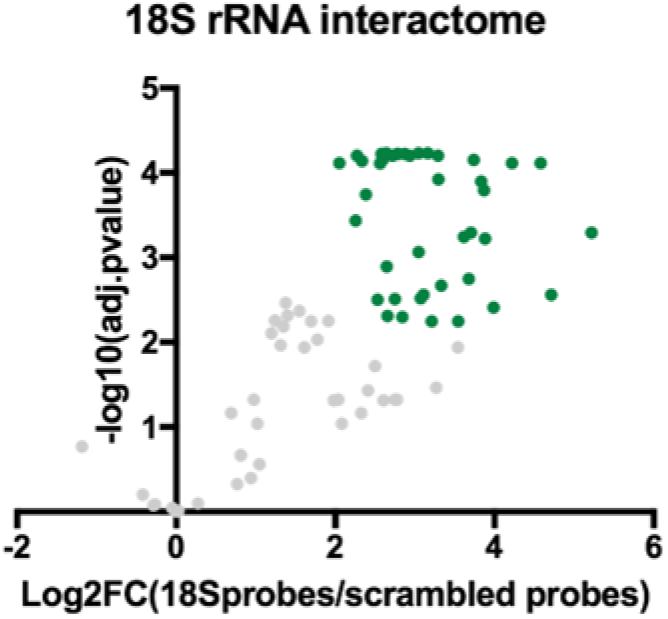
Volcano plot of 5 replicates captured with 18S probes compared with 5 replicates captured with scrambled probes. Green: significantly enriched proteins (adjusted p-value<0.01; |log2FC|>2)

#### Ribosomal proteins of the small subunit

22 of the 33 (66%) proteins of the small subunit (40S formed by the 18S rRNA) were identified (Figure 7A). There are multiple possible explanations for only enriching a subset of all ribosomal proteins of the small subunit. Inefficient UV cross-linking of proteins can occur at RNA-protein interfaces, where the protein interacts with the phosphate backbone. As a result of the preference of amino acid interactions with nucleotide bases, proteins bound to dsRNA stretches are likely to be missed (26),(2). Additionally, smaller proteins have fewer chances of being identified due to a lower number of peptides injected in the mass spectrometer. We indeed observed that mainly larger proteins are identified (Additional file 3). Finally, proteins with only a few direct contacts with the RNA will also result in poor UV cross-linking. For comparison, with RAP-MS (14) 67% of all human small ribosomal proteins were identified, with the use of LNA/DNA mixmer probes (26) only 10%.

#### Ribosomal proteins of the large subunit

10 of the 54 significantly enriched proteins are proteins known to be a part of the large ribosomal subunit (60S formed by 25S rRNA) (Figure 7B). Three of these proteins, RPL19, RPL24 and RPL30, are known to form eukaryotic specific intersubunit bridges to establish the 80S ribosome. RPL19 will interact through its C-terminal α-helical domain with expansion segment 6 of the 18S rRNA (and additionally some small ribosomal subunit proteins: RPS7 and RPS17) forming the eB12 bridge (33),(34). The eB13 bridge is formed by RPL24 interacting with h6,h10 and h44 of 18S rRNA through the linker and α-helix (and additionally a small ribosomal subunit protein: RPS6) (33),(34). RPL30 will form the intersubunit bridge eB9 by interacting with h22 of the 18S rRNA (35). All three are most likely co-purified with the 18S rRNA and not as contamination through their protein-protein interaction with 40S ribosomal proteins (RPS6, RPS7 and RPS17; all enriched in the 18S rRNA interactome). Evidence for this is provided by the described intersubunit interaction eB1b between RPS18 and RPL11. RPS18 is found in our dataset, whereas its protein interactor RPL11 is not (36).

#### Non-ribosomal proteins

22 of the 54 proteins, enriched in the 18S rRNA interactome, are non-ribosomal proteins.

##### Proteins with a (potential) role in rRNA biogenesis

11 of the 22 non-ribosomal proteins are known to play a role in rRNA biogenesis (Figure 7C). All of these are described to be (transient) interaction partners of the rRNA (37),(38),(39),(40),(41),(42),(43),(44),(45),(46).

Five out of the 22 non-ribosomal proteins significantly enriched proteins in the 18S rRNA interactome (Figure 7D) are not yet described to physically interact with 18S rRNA but the literature suggests a potential link with the ribosomal small subunit. In short, PAB1 plays a key role in translation initiation (47). BFR1 and SCP160 often co-purify with polysomes, also suggesting a role in translation (48, 49). SBP1 is known to play a role in translation inhibition of PAB1 by a not yet fully elucidated mechanism (50). Lastly, PUB1 is involved in translation termination through interaction with eRF3, however, this interaction could not be functionally validated, suggesting other mechanisms/interactions to be co-involved (51).

##### Proteins without known ribosome-related association

Six of the 22 non-ribosomal proteins appear to not have a link with the small ribosomal subunit (Figure 7E). These proteins can be either contamination or not yet described to be functional in rRNA biogenesis.

To conclude, 22 of the 54 (quantitative and semi-quantitative) proteins (40.7%) were identified as ribosomal proteins of the small subunit. 10 out of the 54 proteins (18.5%) are identified as ribosomal proteins of the large subunit, of which three (5.5%) are known to directly interact with 18S rRNA. 11 of the 54 proteins (20.4%) are non-ribosomal proteins with a known function in rRNA biogenesis. five out of 54 proteins (9.3%) are proteins with a potential link to rRNA biogenesis and lastly, six out of 54 (11.1%) proteins do not have a known role in rRNA life. In total 75.9% of all significantly enriched proteins of the 18S rRNA interactome appears to be (potential) interactors of the 18S rRNA (Figure 7F). Using RAP-MS (14) 93% of all proteins are potential interactors, with the use of LNA/DNA mixmer probes (26) 64%. Although RAP-MS appears to be somewhat more efficient, we believe that the strength of the protocol we present is its highly cost-effective nature, enabling research labs to perform this type of experiment on a larger scale and with more replication. 24.1% of all enriched proteins are not known yet to interact with 18S rRNA. This underlines the necessity of a proper validation of all targets identified with this type of experiment.

**Figure 7.**
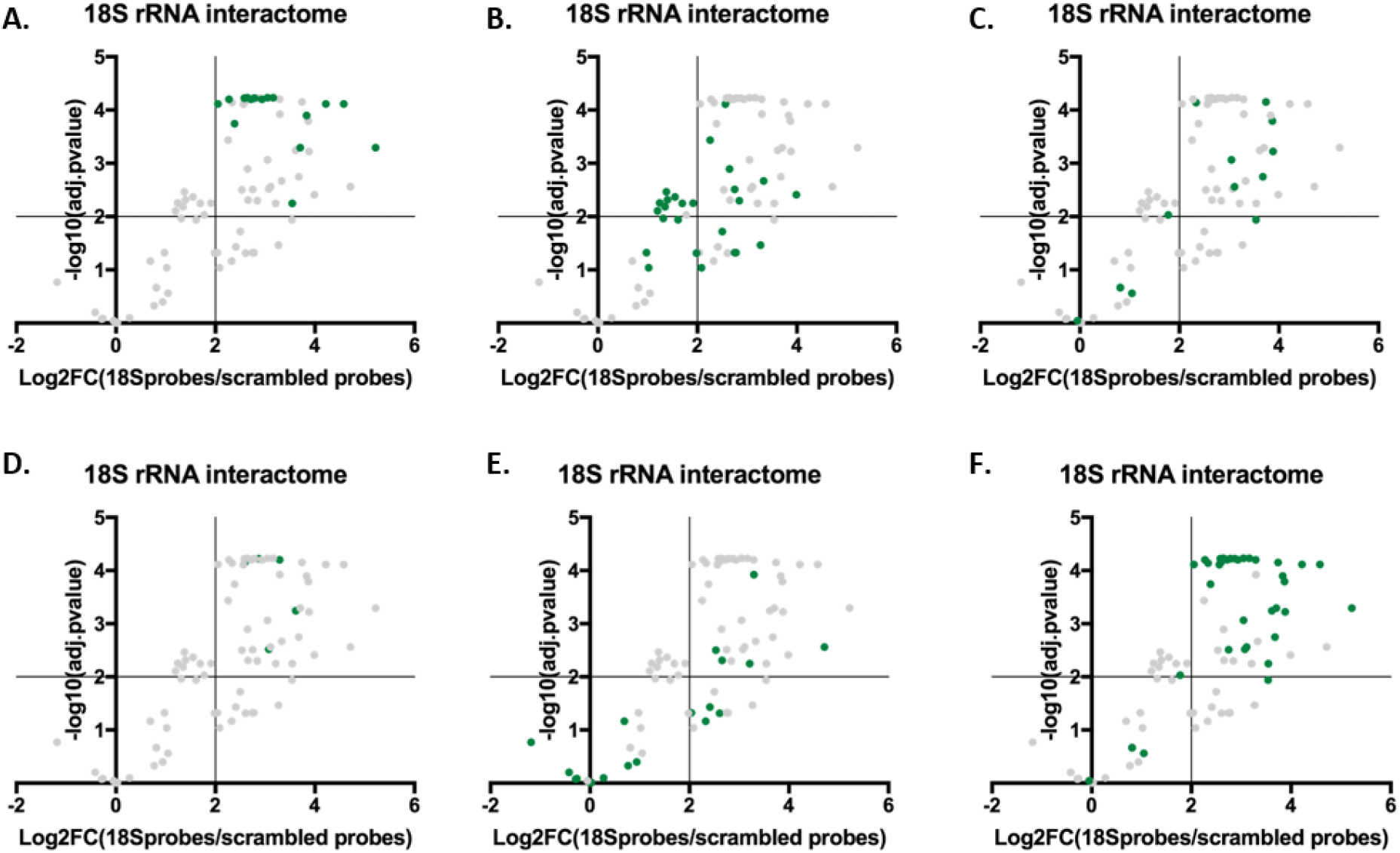
Visualization of different protein groups on volcano plot. A. proteins of the small ribosomal subunit. 1. B. proteins of the large ribosomal subunit C. proteins with a role in rRNA biogenesis D. proteins with a potential role in rRNA biogenesis E. proteins without known ribosome-related association. F. summary of proteins (potentially) interacting with 18S.

#### CryoEM structure

To inspect for biases and correlations, we visualized the significantly enriched ribosomal proteins in the cryoEM structure of this complex. Figure 8A shows the probe distribution along the 18S rRNA. Figure 8B pictures all 22 identified proteins of the small ribosomal subunit. Figure 8C visualizes the ribosomal proteins of the small subunit that were not enriched in our dataset. Aside from size and protein-RNA contact sites, there appears to be a dependency of protein localization contributing to not being identified, as these proteins are grouped. However, we don’t see an immediate explanation for this. Figure 8D shows the enriched ribosomal proteins of the large subunit, with in dark blue the proteins interacting with 18S rRNA. Seven proteins of the large ribosomal subunit are likely to be contaminants of the protocol. When studying p-values, it is clear that these contaminants have larger p-values but remain significant. A more stringent cut-off for example p<0.001 would result in four of the seven ribosomal protein contaminants to become not significant. In addition, four of the six proteins without a described ribosome-related association would be as well labelled as not significant. Only one of the 22 proteins of the small subunit and one intersubunit bridging protein would not be enriched if using this more stringent analysis. Alternatively, instead of using stringent cut-offs, analysing more replicates could contribute to an even more clear discrimination between interactors and contaminants.

**Figure 8.**
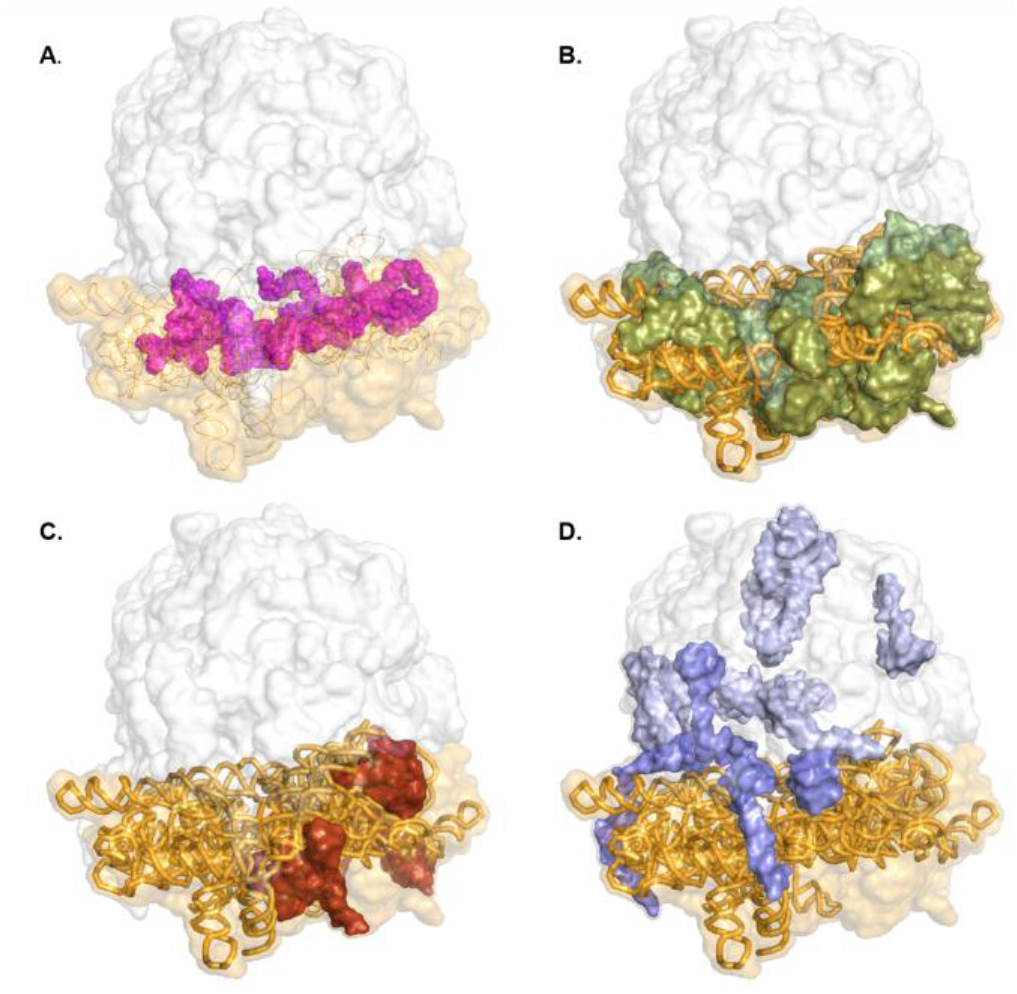
CryoEM structure of the yeast ribosome with in orange: 18S rRNA, white shades: large ribosomal subunit, orange shades: small ribosomal subunit. A. Distribution of the probes along 18S rRNA B. Significantly enriched proteins of the small ribosomal subunit. C. Proteins of the small ribosomal subunit not significantly enriched in our dataset. D. Significantly enriched proteins of the large subunit. In dark blue, the proteins interacting with the 18S rRNA.

### Key optimizations for the establishment of the direct capture protocol

Three main steps, which are not strictly necessary, but enhance the protocol greatly and therefore are strongly recommended were (**1**) the optimization of the ratio probes and beads, resulting in the highest yield and lowest contamination of 25S rRNA. In addition, adding the correct amount of beads is not only cost-effective but will also decrease the peak of unnecessary streptavidin peptides in mass spectrometry. Also contributing to a decreased amount of streptavidin is (**2**) the use of protease-resistant beads (52), which additionally avoids the use of heat or benzonase treatment to elute the proteins. (**3**) Adding formamide to the washing buffers to enhance RNA integrity and capture specificity at reduced temperatures to maintain RNA integrity.

#### Optimal probes/beads ratio

First, the optimal probes/beads ratio compared to the copy number of the target in the input sample was determined. Maximization of the yield (determined as the copy number of 18S rRNA molecules) combined with a minimization of background noise (determined by measuring relative levels of 25S rRNA) for the lowest amount of both probes and beads required was determined. A concentration range of the probe mix was tested with a fixed number of beads (0.5 mg) and input. The minimal amount of probes required was determined to be an excess of 2,000 compared to input. We next investigated whether the reduced yield for the highest amount of probes (an excess of 200,000) was a consequence of the saturation of the streptavidin-coated magnetic beads. As shown in Figure 9, the yield does not increase with an increasing amount of beads disproving this hypothesis. For the minimal amount of probes (excess of 2000) required the amount of beads was minimized as well. The lowest amount of streptavidin-coated magnetic beads required appeared to be 0.250 mg. For this amount, the beads are saturated as further increasing the amount of beads does not result in an increased yield.

**Figure 9.**
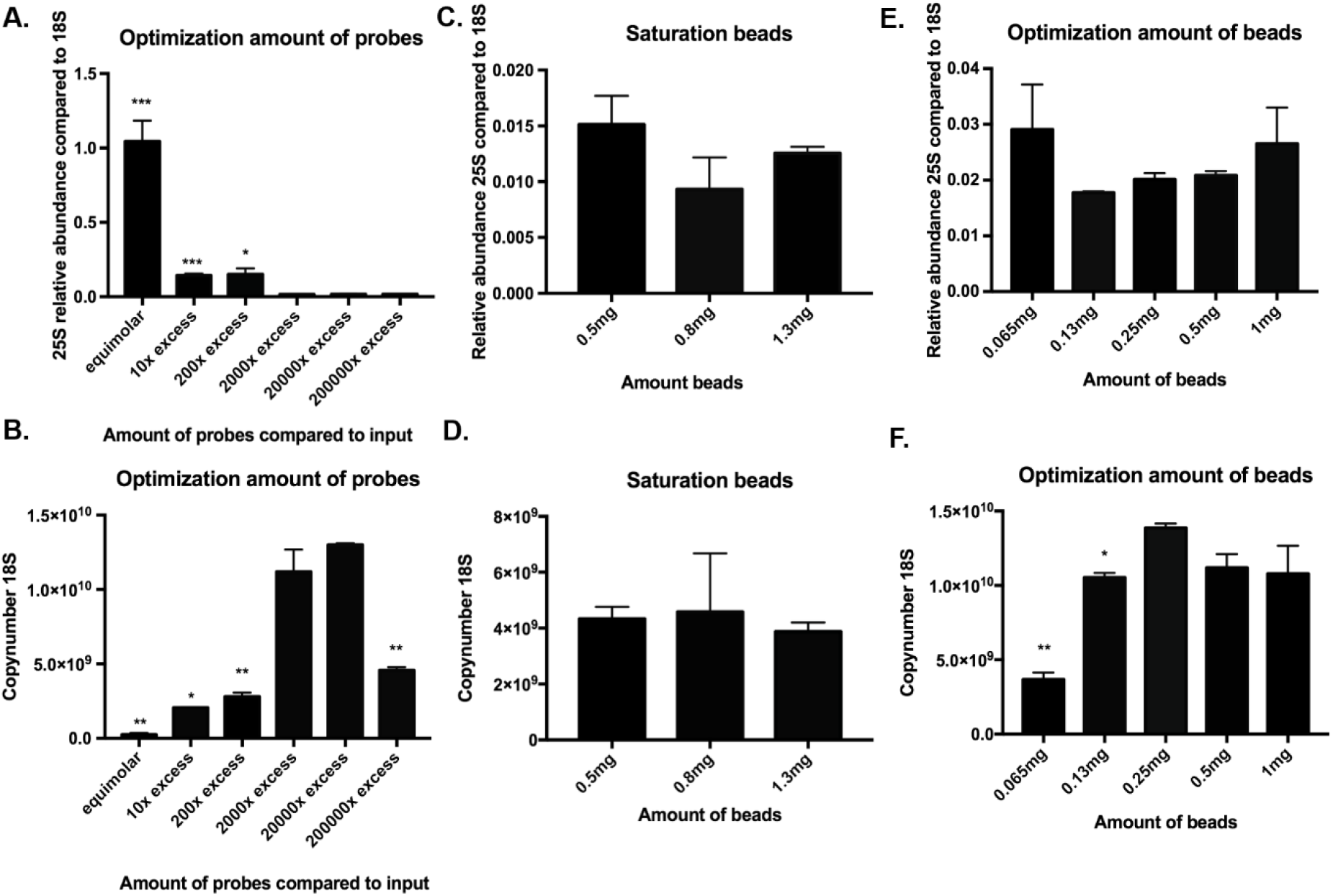
Determining amount of probes and beads required. *** represents a two-tailed p-value<0.001, ** a two-tailed p-value<0.01, * a two-tailed p-value<0.05 A. Relative abundance of 25S compared to 18S for different amount of probes (input 1.5 ×109 and 0.5mg beads). Values are calculated as the amount of 25S after capture divided by the amount of 18S after capture. Error bars represent SEM, n=2. B. Yield after capture for a different amount of probes. (input 1.5 ×109 and 0.5mg beads). Values are calculated by extrapolation on a standard curve of the plasmid PGEM-T_fulllength18S. Error bars represent SEM, n=2. (A-B) Significance is determined with an unpaired t-test compared to 2,000x excess. C. Relative abundance of 25S compared to 18S for different amount of beads (input 1.5 ×109 and 200,000x excess probes) Values are calculated as the amount of 25S after capture divided by the amount of 18S after capture. Error bars represent SEM, n=2. D. Yield after capture for different amount of beads (input 1.5 ×109 and 200,000x excess probes). Values are calculated by extrapolation on a standard curve of the plasmid PGEM-T_fulllength18S. Error bars represent SEM, n=2. (C-D) Significance is determined with an unpaired t-test compared to 0.5mg. E. Relative abundance of 25S compared to 18S for different amount of beads (input 1.5 ×109 and 2,000x excess probes). Values are calculated as the amount of 25S after capture divided by the amount of 18S after capture. Error bars represent SEM, n=2. F. Yield after capture for different amount of beads (input 1.5 ×109 and 2,000x excess probes). Values are calculated by extrapolation on a standard curve of the plasmid PGEM-T_fulllength18S. Error bars represent SEM, n=2. (E-F) Significance is determined with an unpaired t-test compared 0.25mg.

#### Formamide

By using formamide, which destabilizes hydrogen bonds of nucleic acids, the melting temperatures could be lowered by 10-20 ᵒC (0.5ᵒC/%formamide) without losing specificity or yield (Figure 10). Aside from the positive effect of lower temperatures on RNA integrity, formamide also destabilizes RNases (53), which is again beneficial for sample integrity (Figure 11). For the washing steps, the formamide concentration is lowered to 20% for its potential destabilizing effect on biotin-streptavidin interaction. For the last washing step with low salt buffer, formamide is omitted to avoid interference of formamide remnants with the trypsin digestion. Often this type of protocol is performed at lower temperatures, however by determining all temperatures based on the buffer composition and the melting temperatures of the probes, more nonspecific binders will be eluted during washing steps. In addition, secondary structures of the RNA molecule will be denatured in the first step, resulting in more efficient binding of the probes and circumventing the need for a preliminary assay to determine secondary structures such as an RNase H assay (54).

**Figure 10.**
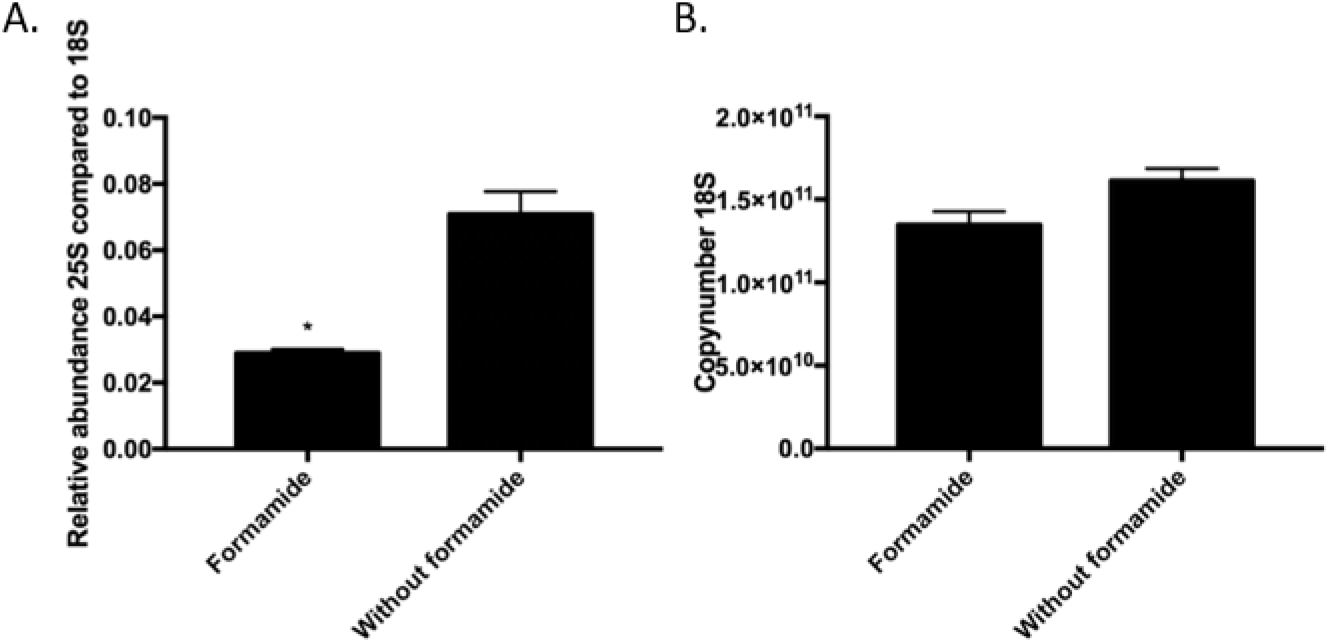
Effect of formamide * represents a two-tailed p-value<0.05 A. Relative abundance of 25S compared to 18S with or w/o formamide. Values are calculated as the amount of 25S after capture divided by the amount of 18S after capture. Error bars represent SEM, n=2. B. Yield after capture with or w/o formamide. Values are calculated by extrapolation on a standard curve of the plasmid PGEM-T_fulllength18S. Error bars represent SEM, n=2. (A-B) Significance is determined with an unpaired t-test compared to 2,000x excess.

**Figure 11.**
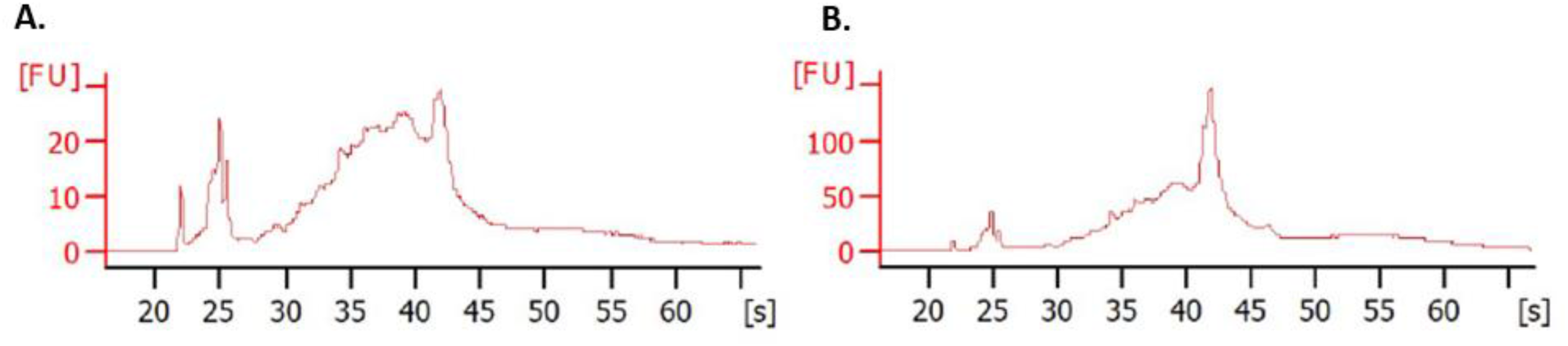
Pico BioAnalyzer (Agilent) A. Capture 18S rRNP w/o formamide B. Capture 18S rRNP with formamide.

### SAPS protocol for multilayer tissues

Next, we tested the applicability of the protocol to multilayer tissues, namely *A. thaliana* mature leaves. The main goal was to establish a successful SAPS experiment for a multicellular organism. We believe that the sole modifications to the procedure are (**1**) the use of an appropriate organism-/tissue-dependent lysis buffer. For *S. cerevisiae*, the lysis buffer described in the TRAPP protocol was used. For *A. thaliana* mature leaves, the plant-specific TRIsure^TM^ lysis buffer was used (figure 12). (**2**) The optimization of the UV cross-linking of multilayer tissues, which is generally challenging due to the inefficient penetration of the light. This can be even more limited by, for example the presence of a cell wall or UV absorbing molecules. Different methods are currently available to introduce covalent links between the RNA (of interest) and its protein interaction partners (18). Despite its low efficiency, we preferred UV light because of its highly specific cross-linking ability. UV254nm was chosen because UV365nm or PAR cross-linking makes use of photoactivatable nucleosides introducing a dependency on compatible sample types which would limit the universal character of the procedure. The penetrability of UV light is highly sample and tissue dependent and dependent on the complexity of the tissue (e.g. multilayer tissue, UV absorbing molecules, presence of a cell wall etc.). We decided to explore an alternative enhanced UV cross-linking procedure to circumvent these tissue dependencies as much as possible. We applied different doses of UV on frozen powdered tissue which, in a way, mimics a cell culture (55).

**Figure 12.**
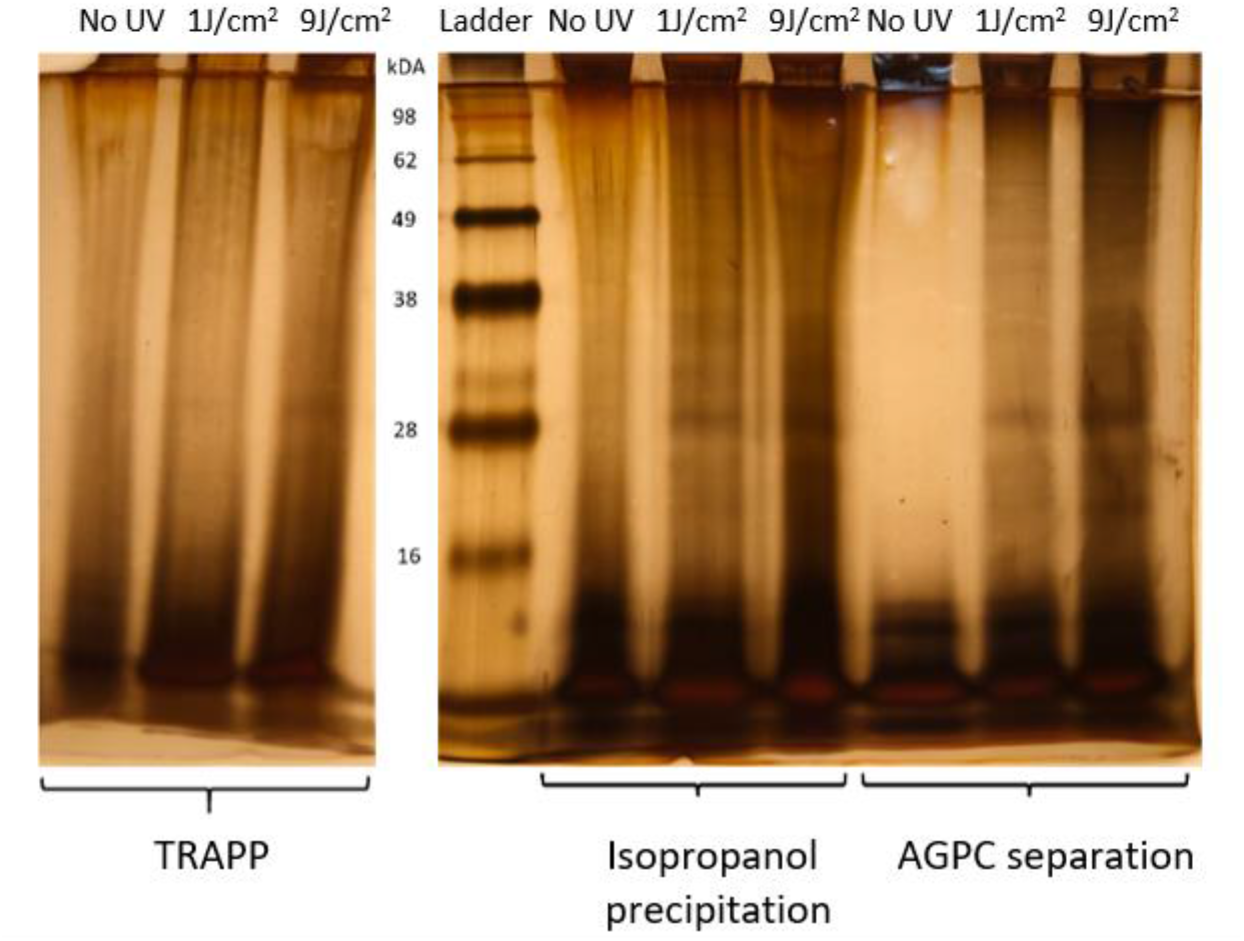
Overview of non-cross-linked and UV cross-linked sample (both 1 and 9 J/cm2) isolate after consecutively TRAPP, Isopropanol precipitation and AGPC separation visualized by a silver stain assay.

### LC-MS/MS analysis study of the RBPome

#### SAPS reveals a confident RBPome for Arabidopsis

To assess the potential of SAPS to purify plant RBPs, a Gene annotation and a Pfam domain analysis have been performed on the general SAPS protein set to verify whether an enrichment in RBP terms could be observed. RNA dependent GO terms are substantially enriched (Figure 13) showing that the SAPS isolation protocol targets the RBPs. We manually compensated for the propagation of GO terms to account for the intrinsic hierarchy of the GO terms. Approximately 58% of the proteins were annotated as RNA-binding. 10% were linked to RNA activity such as catalytic activity acting on RNA, ribonucleoprotein complex binding and translation factor activity. 14% of the proteins could not be linked to any RNA activity and 18% of the gene ID tags could not be linked to a GO term and therefore could not be assigned to one of the above groups. The range of these numbers is comparable to the previous RIC experiments on multiple organisms (56),(57),(58),(59),(60). The proteins of these last two groups could be interesting as previously unknown RNA-interactors. To clarify this, protein-centric approaches such as CLIP could be applied to verify their RNA-binding character. Comparing the sequence and molecular function of newly verified RBPs in these unknown sub-sets could reveal new RNA-binding domains and characteristics of RNA interactors. Additionally, a Pfam study of RNA-binding domains was performed. Approximately, 59% of the SAPS RBPome contain RBDs and 41% domains not linked to RNA (Additional file 4).

**Figure 13.**
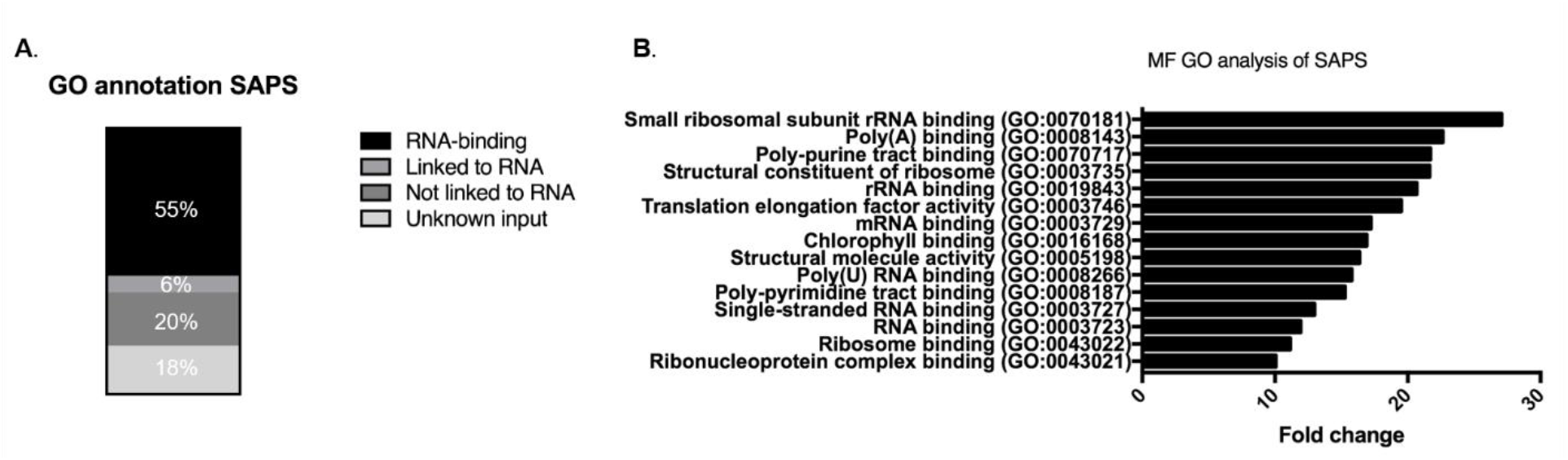
An insight into the RBPome of Arabidopsis thaliana leaf tissue isolated by SAPS. A. The distribution of the RBPome with the GO annotation RNA-binding, linked to RNA, not linked to RNA or proteins whose functions are unknown to date yet. B. The 15 most enriched Molecular functional GO terms represented within the SAPS RBPome clearly linked with RNA-binding terms.

#### Influence of UV cross-linking conditions on the RBPome

Using the above-mentioned specifications, 357 (0,45J), 357 (9J), 493 (P1J) and 526 (P9J) with a total of 709 unique high-fidelity RBP families were identified by the combined UV conditions. Interestingly, most proteins were identified under the conditions using liquid nitrogen flash-frozen ground powder as a source material to perform UV cross-linking on. This observation suggests that cross-linking frozen powder is more efficient as compared to fresh leaf tissue. This is probably due to mimicking monolayer cell cultures, avoiding the numerous obstructions the UV light must pass to reach the RNP complexes. If this interpretation is correct, this approach could be a convenient way to study RNP complexes in vivo in theoretically all kinds of difficult to UV cross-link tissues. There will be no need to compromise on real in vivo studies, by mimicking a setup in protoplasts, cell cultures/suspensions, etiolated plant material, solely because of the tissue type (55).

Additional file 5 shows the distribution of all the identified RBPs between our four conditions. Although the plants used were grown in identical conditions and harvested at the same time, 31% RNA binders unique to one of the conditions could be observed. Similar observations were made by the RBPome studies in multiple organisms summarized by Hentze et al (2018)(2).

A substantial difference in the number of identified RBPs was observed depending on the UV cross-linking condition (Figure 14). Cross-linking of fresh leaf material identified 357 protein families, both for the 0,45J and 9J conditions with an overlap of 55 % between the two conditions. UV cross-linking performed on frozen leaf powder performed better in terms of quantity (P1J identified 493 and P9J 526 protein families to be RBPs) and consistency with a 67% overlap between both conditions. If fresh leaf and frozen powder conditions were compared, an overlap of only 51% is observed. This indicates that the strongest influence on the amount and kind of RBPs as a whole was induced by the sample type (fresh leaves or frozen powder) and to a lesser extent by the UV dose. When a molecular function (MF), biological process (BP) and cellular component (CC) GO-analysis was conducted on both the unique and the overlapping protein IDs of fresh leaf and frozen powder conditions, the following conclusion can be drawn: both the overlapping and unique 84 IDs are highly enriched in MF RNA-binding terms representing the expected RNA-binding character of the protein set (Figure 15).

**Figure 14.**
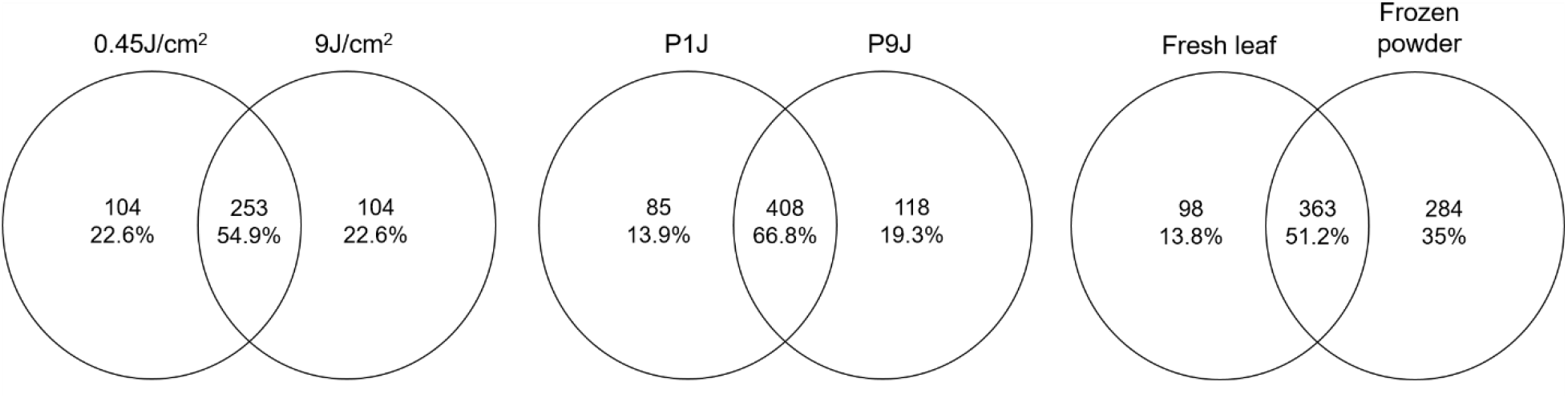
Overlap between the RBPomes of the different sample conditions.

**Figure 15.**
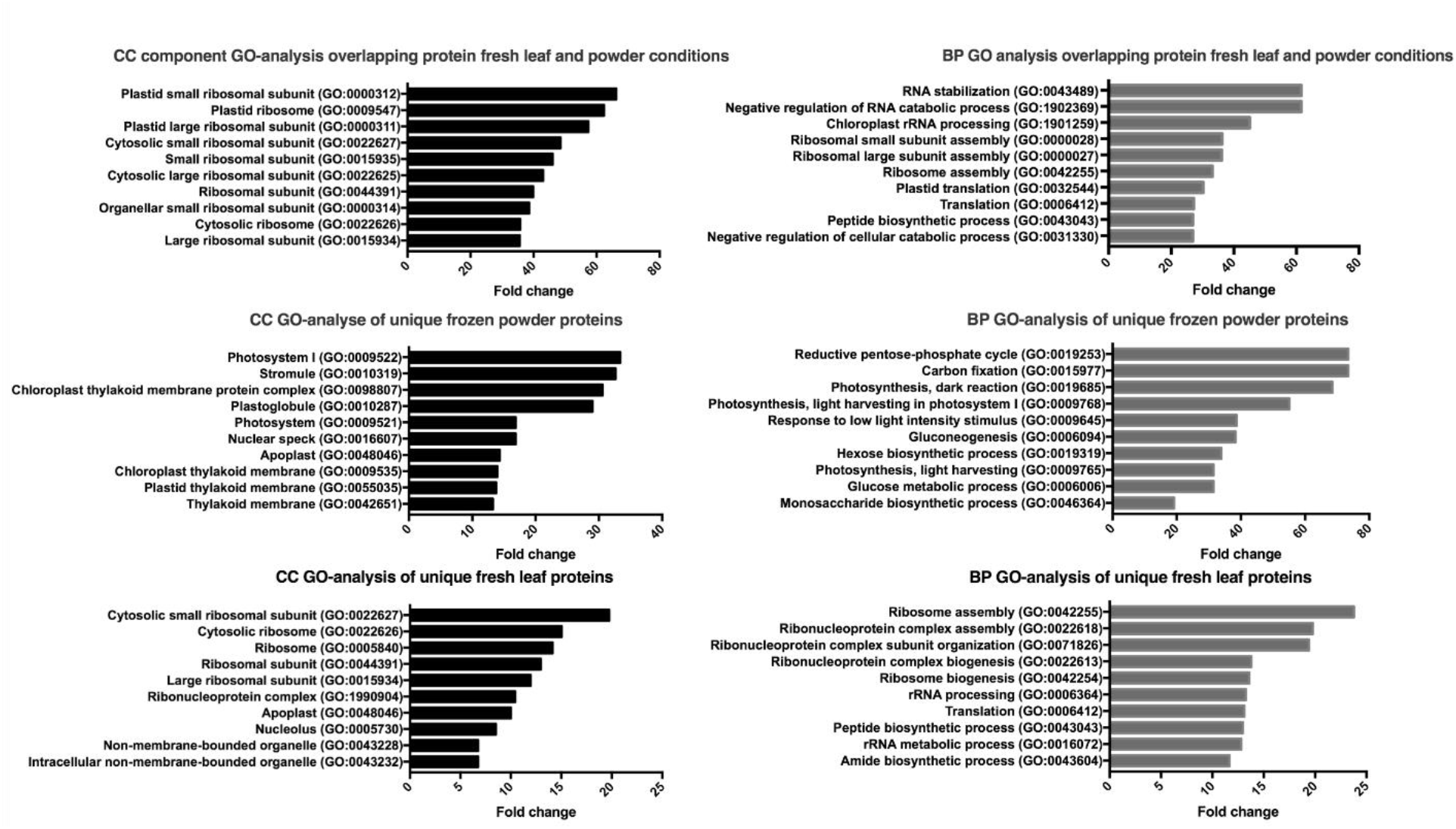
Biological Process (BP) and Cellular Component (CC) GO analysis of the fresh leaf and frozen powder conditions as also their overlap. The 10 highest in terms of enriched fold change are represented for every analysis.

The unique IDs of the frozen powder condition are enriched in BP GO terms linked to expressing photosynthesis, gluconeogenesis and carbon fixation resembling healthy cells building up reserves. The BP terms of the fresh leaf condition are more ribosomal, ribonucleoprotein and translationally orientated lacking these photosynthesis and gluconeogenesis terms. This difference could be explained by the fact that fresh leaf material is still living tissue at the moment of UV cross-linking. The intense UV light could provoke rapid changes upon the RBPome until the samples are flash-frozen. Although cooled on icy water the 0,45J and 9J conditions were exposed to around 2 min and 25 min of extreme UV irradiation during the cross-linking period, respectively. UV stress can induce upregulation of stress-related pathways and downregulation of photosynthesis-related gene expression (61),(62). This perception of UV light as stress is also apparent from the BP terms in the GO analysis of the fresh leaf conditions. This may result in the dynamic changing of the binding characteristics of the RBPome. This UV stress influence could possibly impede interesting RBPome dynamics in the research on the RNA binding character of RBPs upon a biological cue.

## DISCUSSION

### UV cross-linking

The SAPS procedure isolating RNPs was successfully optimized for baker’s yeast and also applied to *A. thaliana.* To overcome the challenge of UV cross-linking multilayer tissue samples instead of cell cultures, liquid nitrogen flash-frozen ground powder mimicking a monolayer cell culture was cross-linked. Together with a more complete set of RBPs, less stress-related proteins were identified when compared with the UV cross-linking of fresh leaves. With UV cross-linking being the main bottleneck for applying SAPS-capture to more complex organisms/tissues compared to yeast, we believe that the introduction of cross-linking flash-frozen ground powder removes this hurdle. The isolation of a confident RBPome for *A. thaliana*, a whole tissue sample with a cell wall, indicates that the translation to other organisms such as mammalian cells will not generate many difficulties. Once SAPS is performed, the RBPome loses its tissue type-dependent character to allow for a universal and streamlined continuation of the RNP isolation procedure. Besides, the dose of UV determines to a lesser extent the amount of RBPs identified. For this reason, we don’t think it is necessary to optimize this step for every organism.

### Silica-based acidic phase separation

For the purification of the complete repertoire of RNPs, the silica-based purification as described by Asencio et al. (2018) (63) and Schepachev et al. (2019) (10) is proven to be a successful strategy when studying the dynamics of the RBPome. However, when the goal is the prepurification for a single RNP isolation, the abundance of non-cross-linked RNA will decrease the RNP/bead ratio. For this reason, we decided to combine the silica-based purification with an AGPC separation. This combination was previously described by Trendel et al. (2019) in the XRNAX protocol (16). However, their protocol starts with an AGPC separation followed by a silica-based separation. When performing the AGPC separation on a more complex sample such as cell lysate, this will result in a less clean interphase potentially trapping unbound RNA. This is not eliminated with the silica-based clean-up when performed afterwards and will as well decrease the RNP/beads ratio by hybridizing with the probes. We chose to reverse this order by performing AGPC separation on a silica-pre-purified sample. This is less time-consuming and generates more soluble interphases only containing RNPs. Alternatively, the interphase could be washed three to four times by repeated AGPC separation to release unbound proteins and RNA molecules, as described by Queiroz et al. (2019) in the OOPS protocol (15). However, this procedure as well requires a subsequent clean-up due to the glycosylated proteins, which share physiochemical properties with RNPs, also co-precipitating on the interphase. This is taken into account by an RNase treatment of the sample and a final AGPC separation. The released RBPs will migrate to the organic phase and by recovering these a pure RBPome is generated. Due to the RNase requirement, this protocol is not suited for subsequent RNP capture.

The difficult nature of cell lysate as a source material due to the presence of RNases/proteinases/secondary metabolites can limit the buffer flexibility and by consequence the applicability of downstream procedures. The SAPS purification will result in a sample of solely RNPs. As mentioned before, the highly uniform nature of the mixture after SAPS, renders this protocol sample type/organism independent by only changing the lysis buffer. As described by Van Ende et al. (2020)(18), the purified sample, lacking lipids, radicals, salts, biotin-containing molecules etc. can be the starting point for many downstream processes, such as studying dynamics of the RBPome, RBD mapping, the study of the protein-bound transcriptome or as described here, the isolation of a specific RNP complex.

### The capture of a specific RNA of interest

We believe that an important strength of this protocol is its cost-effective nature. Due to the low UV cross-linking efficiency (generally 1-5%) (64),(6), a lot of unbound RNA remains present in the cell lysate. These unbound molecules will compete with RNPs to hybridize with the probes and will decrease the RNP/bead ratio drastically. By including SAPS before RNA-capture, only the pool of RNPs remains. The absence of unbound RNA increases the efficiency of the protocol by 95-99% and as a consequence decreases the cost significantly (18). An additional advantage is the removal of the naturally biotin-linked proteins, making the pre-clearance of the cell-lysate with streptavidin-coated beads unnecessary (14). By adding formamide in almost all buffers, the melting temperature could be lowered without losing stringency and thereby avoiding the need for more expensive modified probes (26) or temperature-induced RNA degradation. As a rough estimation, per replicate the total experimental cost (excluding costs for mass spectrometry) is around 60 euro. For comparison, the experimental cost of RAP-MS will be around 2,600 euro per replicate and this is only considering the cost of the beads.

All currently available methods isolate either overexpression targets or highly abundant targets. For low abundant targets, if the material is not limited, our protocol can be easily scaled. The sample will be concentrated during the AGPC separation resulting in workable volumes. A good negative control is of great importance. Examples are scrambled probes, RNase-treated samples, non-cross-linked samples, the capture of another RNP, knock-out samples, etc. Generally, a non-cross-linked control is preferred due to probe or target specific background contamination. However, due to the stringent purification of SAPS before RNA-targeting, these controls appear to be less interesting. We opted for scrambled probes. However, if the experimental set-up allows, this probe specific contamination can be considered when working with a knock-out sample. For lower abundant targets, a combination of negative controls could increase reliability. To increase the probability of detecting low abundant interacting proteins, the background should be as low as possible. An additional ribosomal RNA depletion, oligo dT capture or a double capture (Additional file 6) can increase the detection of the low abundant targets.

## CONCLUSIONS

To conclude, we established a cost-effective, widely applicable protocol that first isolates the whole repertoire of RNPs referred to as SAPS. The isolated RBPome can be the subject of study when investigating the dynamic response of these complexes to environmental or physiological cues or can be the starting point for many downstream processes, such as described here, specific RNP isolation. The SAPS-capture protocol was validated for a well-described RNP, namely 18S rRNP of *S. cerevisiae.* Next, the potential of SAPS-capture to be applied to “difficult to handle” samples was validated by investigating SAPS using *A. thaliana* mature leaves, where we could isolate a confident RBPome. This indicates the potential of the SAPS-capture protocol to be routinely used because it is both tissue-and organism-independent and cost-effective. Future experiments will validate its applicability to more lowly abundant targets. However, the current protocol already illustrates how more abundant targets, such as plant RNA viruses, can be captured from deep tissues, such as phloem. That is an application we are currently pursuing.

## METHODS

### Reagents

UVP crosslinker CL-1000: AnalytikJena, USA, 849-30101-2

TRIsure^TM^: Meridian BIOSCIENCE, Belgium, BIO-38033

Nanodrop spectrophotometer ND-1000: Isogen Life Science, The Netherlands, 6211

DNaseI recombinant, RNase-free: Sigma-Aldrich, Belgium, 4716728001

Pierce^TM^ silver stain kit: ThermoFisher Scientific, Belgium, 24612

Pierce^TM^ BCA protein assay kit: ThermoFisher Scientific, Belgium, 23225

Murine RNase inhibitor: New England BioLabs, The Netherlands, M0314S

Phenylmethanesulfonyl fluoride (PMSF): Sigma-Aldrich, Germany, 78830

Formamide deionized Molecular biology grade: PanReac AppliChem, Germany, A2156.0100

SensiFAST SYBR Hi-ROX Kit: GC Biotech, The Netherlands, BIO-92020

BioAnalyzer 2100: Agilent, Belgium,G2939BA

Pierce^TM^ Trypsin Protease, MS Grade: ThermoFisher Scientific, Belgium, 90057

Streptavidin magnetic beads, New England BioLabs, The Netherlands, S1420S

OMIX C18 pipette tips: Agilent, Belgium, A57003100

Benzonase® nuclease: Sigma-Aldrich, Belgium,70664-3

Phoenix Peptide Cleanup Kit: Preomics, Germany, P.O.00023

Lunatic spectrophotometer: Unchained Labs, USA

Ultimate^TM^ 3000 RSLCnano system: ThermoFisher Scientific, Belgium, ULTIM3000RSLCNANO

Nanospray Flex^TM^ Ion Sources: ThermoFisher Scientific, Belgium, ES071

C18 Reprosil-HD: Dr. Maisch, Germany, r15.b9.

Ultimate^TM^ 3000’s column oven: Thermofisher Scientific, Belgium, 5730.0010

Silica PicoTip emitter: New Objective, USA, FS360-20-10-N-20-C12

µPAC™ HPLC Columns: ThermoFisher Scientific, Belgium, COL-NANO200G1B

Waters nanoEase M/Z HSS T3 Column: Waters Corporation, UK,186008818

### Biological Resources

*Saccharomyces cerevisiae:* S288C

*Arabidopsis thaliana*: Colombia-0

p-GEM®-T vector systems: Promega, The Netherlands, A3600

### Web Sites/Data Base Referencing

Probe design: http://array.iis.sinica.edu.tw/ups/index.php

Genome S288C: https://www.yeastgenome.org/strain/s288c

Scrambled probe design:

https://www.ncbi.nlm.nih.gov/genome/?term=txid12295[Organism:exp]

Blast: https://blast.ncbi.nlm.nih.gov/Blast.cgi

Calculation melting temperatures: https://sourceforge.net/projects/melting/

MaxQuant algorithm (version 1.6.17.0 for *A. thaliana*/ version 2.0.1.0 for *S. cerevisiae*) Protein sequences *A. thaliana*: Swiss-Prot database (database release version of 04_2020) Protein sequences *S. cerevisiae*: Uniprot database (database release version of 11_2020), https://www.uniprot.org/proteomes/UP000002311

Perseus software

R: limma package-moderated t-test

GO analysis: Panther

### Preparation of constructs

Plasmid pGEM-T_fulllength18S was generated by amplifying the full-length 18S by primers PCR1_18SF and PCR2_18SR (Additionally file 7) from *S. cerevisiae* cDNA and inserted into the vector pGEM-T.

### Yeast growth and UV cross-linking

Yeast cells were grown at 30ᵒC under shaking (220 rpm) in YPD medium ((w/v) 1% yeast extract, 2% peptone, and 2% D-glucose). The cells were harvested (10 min, 3000 g) for UV cross-linking (254 nm) at mid-log phase OD 0.5-0.6 from 750 mL of media (roughly 5.5 10^9^ cells). The pellet was resuspended in 200 ml ice-cold cross-linking buffer (65) (25 mM Tris– HCl, pH 7.5; 140 mM NaCl; 1.8 mM MgCl2; and 0.01% NP-40) supplemented with 2% glucose. Per 50 mL, the cells were transferred to a 145/20 mm petri dish and placed on ice in a UVP crosslinker. The cells were irradiated with a dose of 1.2 J/cm^2^. Every 0.4 J/cm^2^, the cells were cooled by swirling for 30 seconds on ice. After cross-linking, the cells were pelleted (5 min,3000 g) and frozen in liquid nitrogen.

### Plant growth and UV cross-linking

Five *Arabidopsis thaliana* plants per pot were grown in day-neutral conditions (12h light, 12h dark) at 20°C with a light intensity of 100 µmol/m^2^/s^2^. Different UV conditions were applied. For fresh leaves, doses of 0,45 J/cm^2^ and 9 J/cm^2^ were applied in a UVP crosslinker. For the dose of 9 J/cm^2^, 10 doses of 0,9 J/cm^2^ were administered with short pauses in between to cool the material. The leaves were placed with the abaxial side upwards on icy water and ice was replenished when thawed. After UV cross-linking, the leaves were patted dry and flash-frozen using liquid nitrogen to preserve the RNA-protein molecular interactions and sample integrity. The frozen powder samples were first ground into powder form, mixed with liquid nitrogen and UV cross-linked in a thin layer of powder/liquid nitrogen mixture. Doses of 1J/cm^2^ (P1J) and 9J/cm^2^ (P9J) were applied. A maximum of 1 J/cm^2^ was applied during each cross-linking run and extra liquid nitrogen was added when necessary to avoid thawing of the samples. Samples were stored at -80°C until further use.

### Silica-based acidic phase separation (SAPS)

The protocol is outlined for 750 mL of yeast cell culture OD 0.5-0.6 or 1 g of plant material but can be easily scaled up/down accordingly.

#### Silica pre-purification of unbound RNA and RNP complexes

In a first step, both unbound RNA and RNPs were purified according to the protocol (total RNA-associated protein purification or TRAPP) described by Shchepachev et al. (2019) (10) with minor modifications. In short these include, (**1**) plant cells were lysed in 10 mL of TRIsure^TM^ supplemented with 10 mM β-mercaptoethanol. The cell lysate was vortexed and incubated for 5 min at RT. (**2**) All centrifugation steps (both for yeast as plant material) to precipitate cell debris were extended to 15 min at 4,750 g. (**3**) During the washing steps, silica beads loaded with the RNA and RNPs were precipitated at 2000 g for 2 min. (**4**) Finally, after elution, the collected eluate was centrifuged for 5 min at maximum speed at 4°C to remove silica powder remnants, which otherwise interfere with the formation of the interphase. The resulting eluate contains both unbound RNA and RNP complexes.

#### DNase treatment (optional), isopropanol treatment and AGPC isolation of RNP complexes

For every 10 µg of RNA (measured with Nanodrop spectrophotometer), 1 U of DNaseI, supplemented with DNase incubation buffer, was used. Half of the DNaseI was added and the sample was incubated for 30 min at 37°C with occasional mixing. Subsequently, the other half was added and incubated for 30 min at 37°C with occasional mixing.

To remove deoxynucleotides and if no DNase treatment was conducted to concentrate the sample, isopropanol precipitation was performed. The eluate was divided into 750 µl per 2 ml tube. 45 µl of 5 M NaCl and 750 µl of ice-cold isopropanol were added. The solution was cooled and stored overnight at -20°C. The RNP and RNA complexes were pelleted by centrifugation at maximum speed for 15 min at 4°C and washed using 1 ml of 70% ethanol.

The pellet was resolubilized into 200 µl of RNase-free water or 10 mM Tris-HCl buffer (pH 7.5) on ice.

To remove both unbound RNA molecules and remaining protein contaminants, 1.2 ml of Trisure^TM^ was added to every 200 µg of RNA equivalent (measured with Nanodrop spectrophotometer) and mixed vigorously to dissolve all the RNP/RNA molecules properly. If a precipitate was still visible, the mixture was heated to 50°C and vortexed till everything was dissolved. 250 µl of chloroform was added, vortexed and incubated for 5 min on a rotating mixer. The samples were centrifuged at maximal speed for 15 min at 4°C to obtain 3 phases. The aqueous phase was removed and the slurry interphase (Additionally file 8) containing the pure RNP complexes was transferred to a new low protein binding tube and dissolved in 200-500 µl 10 mM Tris-HCl RNase-free buffer (pH 7.5). As a quality control, both a silver stain assay and a BCA protein assay were performed. This mixture can be used for the study of the RBPome or as the starting point for the specific RNP-targeting protocol.

### The capture of specific RNP of interest

The final protocol is outlined here. See result section for the optimization procedure.

#### Probe design

Five 60-mer probes with a melting temperature around 70°C were designed to specifically target the RNA of interest (Supplementary table 3). The free software “unique probe selector 2.0” was used. Each DNA oligonucleotide complementary to the RNA sequence of interest was ordered (IDT) with a biotinylated 3’end to enable a capture with streptavidin-coated magnetic beads. For the 18S probes, the RNA sequence of 18S provided by the Saccharomyces Genome Database (SGD) was used. For the scrambled probes, the RNA sequence of Tobacco rattle virus provided by NCBI was used as a template. (These probes were used because of availability in the lab). The scrambled probes were blasted against the genome of *Saccharomyces cerevisiae* to minimize off-targets.

#### RNA-targeting protocol

The protocol is described for one capture. For every replicate, 12 captures were pooled. Generally, one SAPS isolation as described above is sufficient to provide input material for 12 captures (or even more).

1.5 10^9^ copies of the target RNA (determined by absolute RT-qPCR) were mixed with 0.5 mL hybridization buffer (50 mM Tris-HCl pH 7.5, 5 mM EDTA, 500 mM LiCl, 0.2% SDS, 0.1% sodium deoxycholate, 4 M urea) supplemented with 40% deionized formamide, 0.1 mM PMSF, 8 U RNase inhibitor and 0.5 nanomoles of a mixture of all five probes. This mixture was incubated while shaking (450 rpm) at 65°C for 10 min. The temperature was slowly lowered to 45°C, incubated for 5 min and again lowered to 35°C after which the sample was transferred to ice. 0.25 mg of protease-resistant (52) streptavidin coated-magnetic beads, which were previously washed 3 times with wash and bind buffer (20 mM Tris-HCl pH 7.5, 500 mM LiCl, 1 mM EDTA), were added together with 0.5 mL hybridization buffer. The probe-RNP complexes were incubated together with the beads for 2 hours at 50°C while shaking (450 rpm). Probe-RNP complexes bound to the beads were then washed for 3 min at 60°C with wash and bind buffer supplemented with 20% deionized formamide. This step was performed 2 times. Afterwards, the beads were washed for 3 min at 55°C with low salt buffer (20 mM Tris-HCl pH 7.5, 150 mM LiCl, 1 mM EDTA) supplemented with 20% deionized formamide. The mixture is transferred to a clean low binding tube and a final wash for r 3 min at 55°C with low salt buffer was conducted. 90 µg beads were removed after the final wash and eluted in 5 µl elution buffer (10 mM Tris-HCl pH 7.5) for 3 min at 95°C for quality control using RT-qPCR and RNA pico BioAnalyzer. The remaining beads were resolved in 150 µl trypsin digestion buffer (20 mM Tris-HCl pH 8.0, 2 mM CaCl2) and incubated for 4 hours with 1 µg trypsin at 37°C. Beads were removed, another 1 µg of trypsin was added and proteins were further digested overnight at 37°C. Peptides were purified on Omix C18 tips and dried completely in a rotary evaporator. All used binding and washing temperatures were calculated using the free software “MELTING”.

#### Quality control: RT-qPCR and BioAnalyzer

To determine the purity of the specific RNP samples/specificity of the RNA-targeting protocol, both a RT-qPCR and a BioAnalyzer assay were performed to check abundance of non-target genes after the capture according to the protocol of the manufacturer. A standard volume of 7.5 µl was used in the 10 µl reverse transcription reaction volume. The RNA integrity number (BioAnalyzer), which is based on the 25S/18S ratio, could not be determined due to the absence of 25S after the capture. The BioAnalyzer assay was solely performed to check the presence of abundant RNA contaminants following the manufacturing protocol of the RNA 6000 Pico Chip (Agilent).

### Sample preparation for mass spectrometry study of the RBPome in A. thaliana

For mass spectrometry sample preparation, 2 g of plant material per replicate were used.

The RNA part of the RNP molecules was degraded using Benzonase 99% pure. The RNP mixture was heated to 80°C to dissolve liquid-liquid phase separation complexes that can occur and could impede the Benzonase cleaving efficiency. The samples were cooled to 37°C and 12 U of Benzonase supplemented with 1 mM MgCl_2_ was added. After the Benzonase digest, the SP3 method (66) was used to digest the proteins into peptides and to desalt the samples following the published protocol scaled to our volumes. The peptides were eluted in 100 µl of 100 mM ammonium bicarbonate (pH 8) and sent on dry ice to the mass spectrometry Facility Core of the University of Ghent for further processing. For each experimental condition of the plant samples, part of one replicate was pre-ran on the mass spectrometry set-up as a trial. The presence of a polymer resulting in clogging of the machine was observed. The samples were further purified using the Phoenix peptide clean-up kit successfully removing the polymer.

### LC-MS/MS analysis

Peptides of the 18S rRNA interactome were re-dissolved in 20 µl loading solvent A (0.1% trifluoroacetic acid in water/acetonitrile (ACN) (98:2, v/v)) of which 2 µl was injected for LC-MS/MS analysis on an Ultimate 3000 RSLCnano system in-line connected to a Q Exactive HF mass spectrometer. Trapping was performed at 10 μl/min for 4 min in loading solvent A on a 20 mm trapping column (made in-house, 100 μm internal diameter (I.D.), 5 μm beads, C18 Reprosil-HD). The peptides were separated on a 250 mm Waters nanoEase M/Z HSS T3 Column, 100Å, 1.8 µm, 75 µm inner diameter kept at a constant temperature of 45°C. Peptides were eluted by a non-linear gradient starting at 1% MS solvent B reaching 33% MS solvent B (0.1% FA in water/acetonitrile (2:8, v/v)) in 63 min, 55% MS solvent B (0.1% FA in water/acetonitrile (2:8, v/v)) in 87 min, 99% MS solvent B in 90 min followed by a 10-minute wash at 99% MS solvent B and re-equilibration with MS solvent A (0.1% FA in water). The mass spectrometer was operated in data-dependent mode, automatically switching between MS and MS/MS acquisition for the 12 most abundant ion peaks per MS spectrum. Full-scan MS spectra (375-1500 m/z) were acquired at a resolution of 60,000 in the Orbitrap analyzer after accumulation to a target value of 3,000,000. The 12 most intense ions above a threshold value of 15,000 were isolated with a width of 1.5 m/z for fragmentation at a normalized collision energy of 30% after filling the trap at a target value of 100,000 for maximum 80 ms. MS/MS spectra (200-2000 m/z) were acquired at a resolution of 15,000 in the Orbitrap analyzer.

Purified peptides of the plant sample for shotgun analysis were re-dissolved in 20 µl solvent A (0.1% TFA in water/ACN (98:2, v/v) and peptide concentration was determined by measuring on a Lunatic spectrophotometer. 2 µg (*A. thaliana*) of each sample was injected for LC-MS/MS analysis on an Ultimate 3000 RSLCnano system in-line connected to a Q Exactive HF mass spectrometer equipped with a Nanospray Flex Ion Source. Trapping was performed at 10 μl/min for 4 min in solvent A on a 20 mm trapping column (made in-house, 100 μm internal diameter (I.D.), 5 μm beads, C18 Reprosil-HD) and the plant sample was loaded on a 200 cm long micro-pillar array column with a C18-endcapped functionality mounted in the Ultimate 3000’s column oven at 50°C. For proper ionization, a fused silica PicoTip emitter (10 µm I.D.) was connected to the µPAC™ outlet union and a grounded connection was provided to this union. Peptides were eluted by a non-linear increase from 1 to 55% MS solvent B (0.1% FA in water/ACN (2:8, v/v)) over 145 min, first at a flow rate of 750 nl/min, then at 300 nl/min, followed by a 15 min wash reaching 99% MS solvent B and re-equilibration with MS solvent A (0.1% FA in water). The mass spectrometer was operated in data-dependent mode, automatically switching between MS and MS/MS acquisition for the 16 most abundant ion peaks per MS spectrum. Full-scan MS spectra (375-1,500 m/z) were acquired at a resolution of 60,000 in the Orbitrap analyzer after accumulation to a target value of 3E6. The 16 most intense ions above a threshold value of 1.3E4 (minimum AGC of 1E3) were isolated for fragmentation at a normalized collision energy of 28%. The C-trap was filled at a target value of 100,000 for maximum 80 ms and the MS/MS spectra (200-2,000 m/z) were acquired at a resolution of 15,000 in the Orbitrap analyzer with a fixed first mass of 145 m/z. Only peptides with charge states ranging from +2 to +6 were included for fragmentation and the dynamic exclusion was set to 12 s.

### Identification and quantification of proteins

LC-MS/MS runs of all samples were searched together using the MaxQuant algorithm with mainly default search settings, including a false discovery rate set at 1% on PSM, peptide and protein level. Spectra were searched for *Saccharomyces cerevisiae* against the *Saccharomyces cerevisiae* protein sequences in the Uniprot database containing 6,049 sequences and for *A. thaliana* against the *Arabidopsis* protein sequences in the Swiss-Prot database, containing 39,359 sequences. The mass tolerance for precursor and fragment ions was set to 4.5 and 20 ppm, respectively, during the main search. Enzyme specificity was set as C-terminal to arginine and lysine, also allowing cleavage at proline bonds with a maximum of) two (*S. cerevisiae*)/three (*A. thaliana)* missed cleavages. Variable modifications were set to oxidation of methionine residues and acetylation of protein N-termini, while carbamidomethylation of cysteine residues was set as fixed modification for *A. thaliana* samples. Matching between runs was enabled with a matching time window of 0.7 min and an alignment time window of 20 min. Only proteins with at least one unique or razor peptide were retained. Proteins were quantified by the MaxLFQ algorithm integrated in the MaxQuant software. A minimum ratio count of two unique or razor peptides was required for quantification.

To compare protein intensities in the 18S probes and scrambled probes samples, statistical testing for differences between the two group means was performed, using the R-package Limma (moderated t test). Missing protein intensity values were imputed by randomly sampling from a normal distribution centered around each sample’s noise level. Statistical significance for differential regulation was set at adjusted p-value < 0.01 and |log2FC| = 2. Since our to compare datasets have a large difference in protein intensities (scrambled group should be theoretically lacking proteins) iBAQ intensities were chosen over MaxLFQ intensities for quantification. To appoint proteins to be part of the interactome both a semi-quantitative as a quantitative method, were used. If proteins were not detected in any of the non-cross-linked samples but present in 4 of the 5 replicates of the condition this protein was appointed to be an interaction partner of 18S rRNA in a semi-quantitative manner.

Further data analysis of the shotgun results of the study of the RBPome of *A. thaliana* was performed with the Perseus software and Limma package (moderated T-test) in R. Since the datasets we want to compare have a large difference in protein intensities (control group should be theoretically lacking proteins) iBAQ intensities were chosen over MaxLFQ intensities for quantification. To appoint proteins to be part of the RBPome both a semi-quantitative as a quantitative method, was used. If proteins were not detected in any of the non-cross-linked samples but present in 3 of the 5 replicates of the condition this protein was appointed to be an RBP in a semi-quantitative manner (67). Proteins with an iBAQ value in both the non-cross-linked control and cross-linked conditions a quantitative method was applied. Using Perseus, proteins yielding minimal 3 iBAQ values per 5 replicates were selected, log-transformed and the missing values were imputed with values drawn from a normal distribution. This modified dataset was used to perform the moderated t-test implemented in the R/Bioconductor package Limma. The p-values were corrected for multiple testing using the Benjamini-Hochberg test to calculate the adjusted p-value. Proteins with an adjusted p-value < 0.01 and a |log2FC| (CL/No-UV) > 1.5 were appointed to be true RNA binding proteins.

### CryoEM structures of yeast ribosome

In order to visualise the proteins identified using our here presented approach we used PDB entry: 3JJ7. The images were rendered using pymol. Every chain within the structure was inspected and labelled as being significantly enriched or not. The non-ribosomal Guanine nucleotide-binding protein subunit beta-like protein was removed during visualisation.

### Gene Ontology (GO) enrichment analysis for the study of the RBPome of A. thaliana

Protein IDs of the identified RBPs were used to perform GO-analysis using Panther. The reference *Arabidopsis* proteome was used to calculate enrichment applying the Fisher’s exact test and Bonferroni corrected p-values were used to account for multiple testing. GO-analysis was performed for both Biological process (BP), Molecular function (MF) and cellular component (CC) and compared between the different conditions or subsets within the unique and overlapping protein IDs between different conditions.

## DECLARATIONS

### Ethics approval and consent to participate

Not applicable

### Consent for publication

Not applicable

### Availability of data and materials

The datasets generated and/or analysed during the current study are available in the PRIDE repository, https://www.ebi.ac.uk/pride/archive/

For the 18S rRNA interactome, the mass spectrometry proteomics data have been deposited to the ProteomeXchange Consortium via the PRIDE [1] partner repository with the dataset identifier PXD031573.

**Password:** nHKrsXU8

For the RBPome of *A. thaliana*, the mass spectrometry proteomics data have been deposited to the ProteomeXchange Consortium via the PRIDE [1] partner repository with the dataset identifier PXD031578.

**Password:** bfOD8ODc

### Competing interests

The authors declare that they have no competing interests.

### Funding

This work was supported by het Fonds Wetenschappelijk Onderzoek PhD fellowship strategic basic research. [grant numbers 1S40720N, 1S06517N].

### Authors’ contributions

S.B., R.V.E. and K.G. conceived and designed the study. S.B. and R.V.E. performed the experiments. A.V. visualized the cryoEM structures. S.B, R.V.E. and K.G. wrote the paper. All authors read and approved the final manuscript.

### Authors’ information

The authors wish it to be known that, in their opinion, the first 2 authors should be regarded as joint First Authors.

## Acknowledgements

Not applicable

## Notes

### Competing Interest Statement

The authors have declared no competing interest.

