## additional files for "A widely applicable and cost-effective method for general and specific RNA-protein complex isolation"

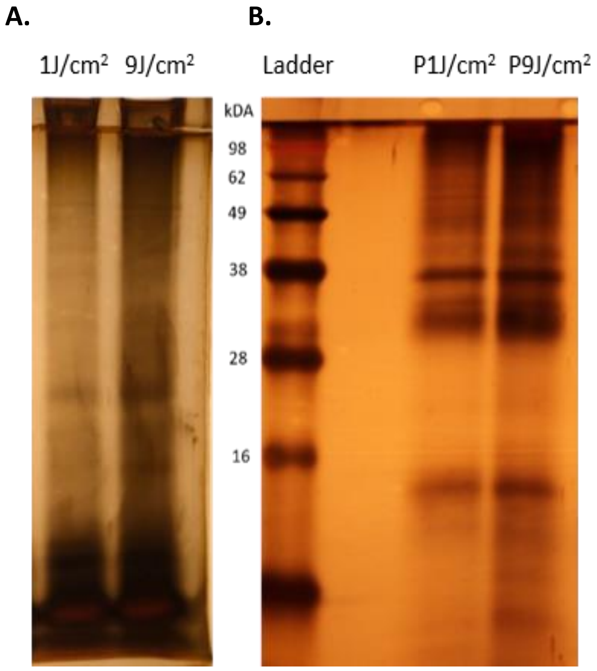


Additional file 1. A. RNase treatment before visualization on silver stain B. Benzonase treatment before visualisation on silver stain.

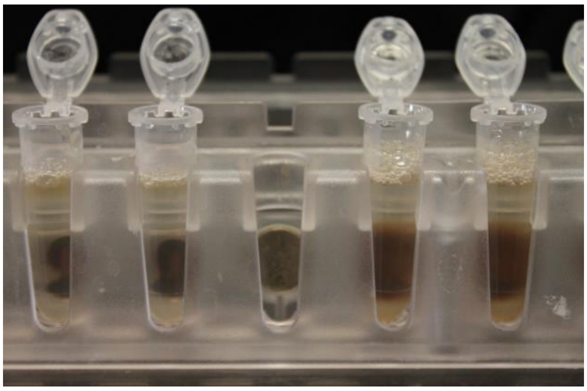

Additional file 2. **Difference in beads dispersion against the magnet.** Left tubes: capture with scrambled probes. Right tubes: capture with 18S probes.

Additional file 3. Small ribosomal proteins sorted by length (**Bolt**: identified by MS)

| Protein | Length (aa) |
|---|---|
| RPS29A, RPS29B | 56 |
| RPS30A, RPS30B | 63 |
| RPS28A, RPS28B | 67 |

| | |
|---|---|
| RPS27A, RPS27B | 82 |
| RPS21A, RPS21B | 87 |
| **RPS10A, RPS10B** | **105** |
| **RPS25A;RPS25B** | **108** |
| **RPS26B;RPS26A** | **119** |
| **RPS20** | **121** |
| RPS22 | 130 |
| **RPS24B;RPS24A** | **135** |
| **RPS17B;RPS17A** | **136** |
| RPS14 | 137 |
| RPS15 | 142 |
| **RPS16A, RPS16B** | **143** |
| RPS12 | 143 |
| **RPS19B;RPS19A** | **144** |
| **RPS23A, RPS23B** | **145** |
| **RPS18B;RPS18A** | **146** |
| **RPS13** | **151** |
| RPS31;RPL40B;RPL40A;UBI4 | 152 |
| **RPS11B;RPS11A** | **156** |
| **RPS7A** | **190** |
| **RPS9B;RPS9A** | **197** |
| **RPS8B;RPS8A** | **200** |
| **RPS5** | **225** |
| **RPS6B;RPS6A** | **236** |
| **RPS3** | **240** |
| **RPS0B;RPS0A** | **252** |
| **RPS2** | **254** |
| **RPS1B** | **255** |
| **RPS4B;RPS4A** | **261** |

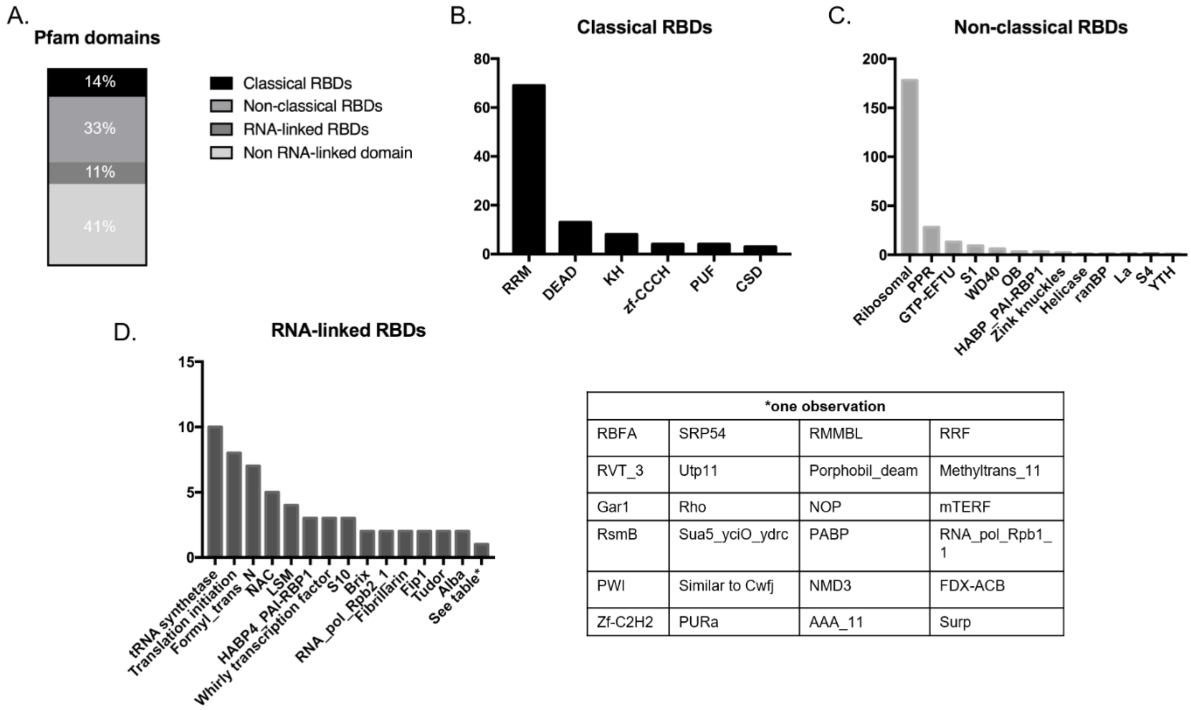


| *one observation | | | |
|---|---|---|---|
| RBFA | SRP54 | RMMBL | RRF |
| RVT_3 | Utp11 | Porphobil_deam | Methyltrans_11 |
| Gar1 | Rho | NOP | mTERF |
| RsmB | Sua5_yciO_ydrc | PABP | RNA_pol_Rpb1_1 |
| PWI | Similar to Cwfj | NMD3 | FDX-ACB |
| Zf-C2H2 | PURa | AAA_11 | Surp |

Additional file 4. **Study of the RNA-binding Domains (RBDs) of the SAPS isolated A. thaliana leaf RBPome.** A. The distribution of proteins harboring classical B. non-classical C. RNA-linked binding domains D. and domains not linked to RNA based on a Pfam annotation.

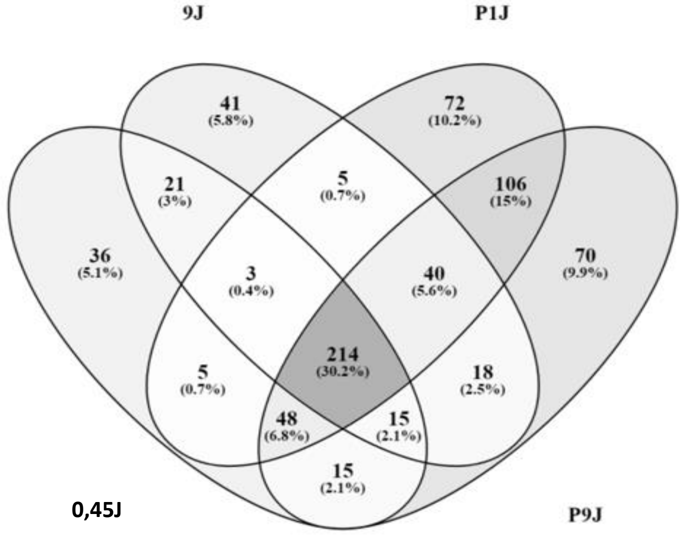


Additional file 5. Comparison of occurring RBPs per UV cross-linking condition. The UV dose has been displayed in the figure where P stands for inducing UV light on frozen powder tissue instead of fresh leaf tissue.

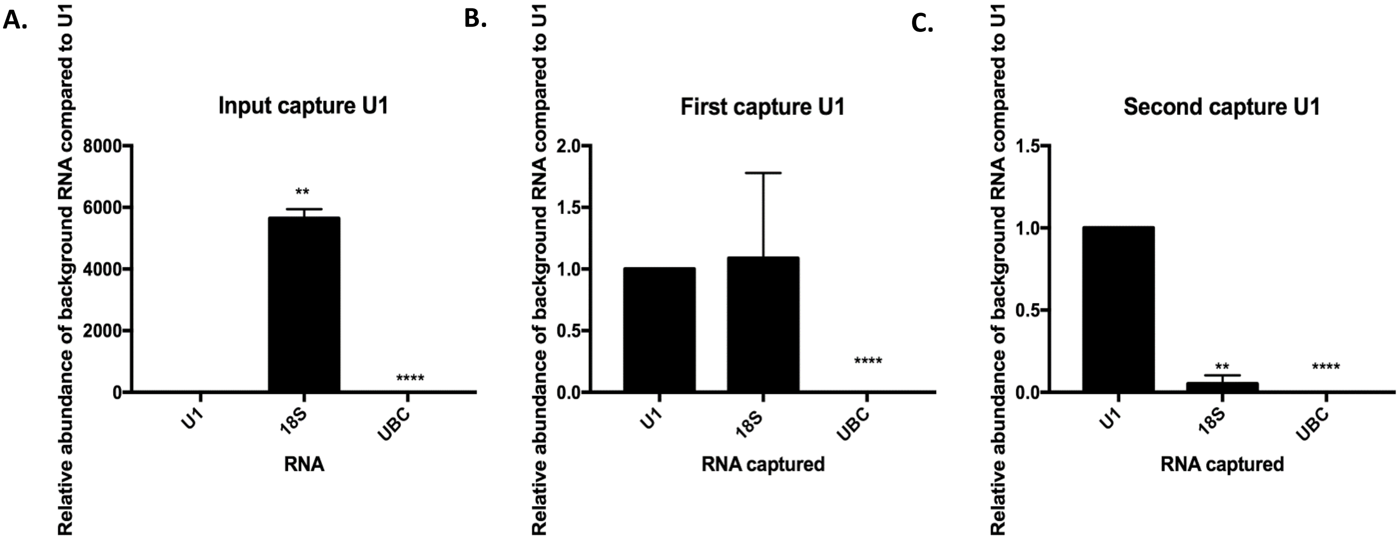


Additional file 6. **Capture of U1 in *S. cerevisiae*** **** represents a two-tailed p-value<0,0001, ** a two-tailed p-value<0,01 A. Relative abundance of 18S/UBC compared to U1 after SAPS. Error bars represent SEM, n=2. B. Relative abundance of 18S/UBC compared to U1 after capture of U1 . Error bars represent SEM, n=2. C. Relative abundance of 18S/UBC compared to U1 after a second capture of U1 . Error bars represent SEM, n=2.

Additional file 7. PCR and RT-qPCR primers

| | |
|---|---|
| **PCR primers** | |
| PCR1_18sF | 5’-aataagggttcgattccggag-3’ |
| PCR2_18sR | 5’-aaagggcagggacgtaatc-3’ |
| **qPCR primers** | |
| qPCR1_18sF | 5’-cctttccttctggctaaccttgagtccttg-3’ |
| qPCR2_18sR | 5’-cgagcaatacgcctgctttgaacactc-3’ |
| qPCR3_25sF | 5’-cccactgtccctatctactatc-3’ |
| qPCR4_25SR | 5’-gctcaacagggtcttctttc-3’ |
| qPCR5_taf10F | 5’-atattccaggatcaggtcttccgtagc-3’ |
| qPCR6_taf10R | 5’-gtagtcttctcattctgttgatgttgttgttg-3’ |

Additional file 8. Probes for RNP-targeting of 18S rRNA

| **18S probes** |
|---|
| 5’tacttagacatgcatggcttaatctttgagacaagcatatgactactggcaggatcaacc /3BioTEG/3’ |
| 5’cggtactagcgacgggcggtgtgtacaaagggcagggacgtaatcaacgcaagctgatga/3BioTEG/3’ |
| 5’ ttaggattgggtaatttgcgcgcctgctgccttccttggatgtggtagccgtttctcagg /3BioTEG/3’ |
| 5’actcgctggctccgtcagtgtagcgcgcgtgcggcccagaacgtctaagggcatcacaga/3BioTEG/3’ |
| 5’ttaactgcaacaactttaatatacgctattggagctggaattaccgcggctgctggcacc /3BioTEG/3’ |
| **Scrambled probes** |
| 5’gtttcaacacgtttacgacacttttcaaggtgactacggccacaacaattatacacatca/3BioTeg/3’ |
| 5’tattagcgacacttcgatactgaaagggtatgtaccgtcacagaccggaacgccctcaga/3BioTeg/3’ |
| 5’cagtccgtgcgggtaattctgacctccagttcgaataactcctctcgaatagtcacgata/3BioTeg/3’ |
| 5’ttcggtagtactcttgcgatgatagagacctcctcggaactccagctatcctgtagatcg/3BioTeg/3’ |
| 5’tgagtgtgcgacacctcccaaaggaaggccgcccactcatctagcccaaataacctgtaa/3BioTeg/3’ |
